# Type I Interferon Limits Viral Dissemination-Driven Clinical Heterogeneity in a Native Murine Betacoronavirus Model of COVID-19

**DOI:** 10.1101/2020.09.11.294231

**Authors:** Hua Qing, Lokesh Sharma, Brandon K. Hilliard, Xiaohua Peng, Anush Swaminathan, Justin Tian, Kavita Israni-Winger, Cuiling Zhang, Delva Leão, Seungjin Ryu, Victoria Habet, Lin Wang, Xuefei Tian, Yina Ma, Shuta Ishibe, Lawrence H. Young, Sergei Kotenko, Susan Compton, Carmen J. Booth, Aaron M. Ring, Vishwa Deep Dixit, Craig B. Wilen, João P. Pereira, Charles S. Dela Cruz, Andrew Wang

**Author notes:** These authors contributed equally to this work (Hua Qing, Lokesh Sharma, Charles S. Dela Cruz, Andrew Wang).

## Abstract

Emerging clinical data demonstrates that COVID-19, the disease caused by SARS-CoV2, is a syndrome that variably affects nearly every organ system. Indeed, the clinical heterogeneity of COVID-19 ranges from relatively asymptomatic to severe disease with death resultant from multiple constellations of organ failures. In addition to genetics and host characteristics, it is likely that viral dissemination is a key determinant of disease manifestation. Given the complexity of disease expression, one major limitation in current animal models is the ability to capture this clinical heterogeneity due to technical limitations related to murinizing SARS-CoV2 or humanizing mice to render susceptible to infection. Here we describe a murine model of COVID-19 using respiratory infection with the native mouse betacoronavirus MHV-A59. We find that whereas high viral inoculums uniformly led to hypoxemic respiratory failure and death, lethal dose 50% (LD_50_) inoculums led to a recapitulation of most hallmark clinical features of COVID-19, including lymphocytopenias, heart and liver damage, and autonomic dysfunction. We find that extrapulmonary manifestations are due to viral metastasis and identify a critical role for type-I but not type-III interferons in preventing systemic viral dissemination. Early, but not late treatment with intrapulmonary type-I interferon, as well as convalescent serum, provided significant protection from lethality by limiting viral dissemination. We thus establish a Biosafety Level II model that may be a useful addition to the current pre-clinical animal models of COVID-19 for understanding disease pathogenesis and facilitating therapeutic development for human translation.

## Introduction

COVID-19, the disease caused by SARS-CoV-2, results in a wide range of clinical phenotypes ranging from asymptomatic to death with manifestations that have been described in virtually every organ system (Gupta et al., 2020; Zheng et al., 2020). One major limitation in developing effective therapies is in the lack of animal models that recapitulate mechanisms of cell and tissue injury and that capture the wide clinical heterogeneity observed in humans. Indeed, widespread access in academic and biopharmaceutical institutions to animal models that can recapitulate the myriad of clinical expressions seen in patients are essential to characterize the pathophysiology of this novel coronavirus infection to maximize therapeutic development. Unfortunately, few species are naturally receptive to infection by SARS-CoV-2. These include model species that are difficult to maintain and genetically modify and include non-human primates (Munster et al., 2020), ferrets (Kim et al., 2020) and hamsters (Chan et al., 2020); moreover, inoculation of SARS-CoV-2 strains isolated from clinical patients does not produce the full clinical heterogeneity observed in COVID-19, limiting their utility as preclinical models.

Mice are the most accessible and versatile model species for preclinical development. However, they are resistant to human SARS-CoV-2 infection due to the low affinity of the mouse ACE2 receptor for the Spike protein. Thus, several groups have recently reported strategies that allow mice to be receptive to SARS-CoV-2 infection either by transgenic expression of human ACE2 (hACE2), expressing hACE2 using adenovirus technology, or mouse-adapting SARS-CoV-2 (Bao et al., 2020; Gu et al., 2020; Hassan et al., 2020; Israelow et al., 2020; Sun et al., 2020a; Sun et al., 2020b). One common characteristic in all of these mouse models is the fairly mild disease expression and limited viral dissemination, which is either a result of the design of the model system (respiratory transduction with adenovirus) or a limitation of the system (transgenic models), precluding the ability to study the full spectrum of disease observed in COVID-19 patients. Indeed, viral dissemination is emerging as a key driver in the pathogenesis of COVID-19 underlying its wide clinical heterogeneity (Puelles et al., 2020). It is likely that in order to develop COVID-19-like disease in animals expressing human ACE2 throughout the body, SARS-CoV-2 will have to be further “murinized” through serial passaging, as was observed in MERS-CoV animal models (Li et al., 2017). In addition to humanizing the mouse with transgenic expression of human DPP4, it was also necessary to murinize the native MERS-CoV in order to develop an animal model of human disease (Li et al., 2017). However, the resultant mouse-adapted MERS-CoV 22 contained over 22 novel mutations, including several within the spike protein, undermining, to some extent, the rationale for this experimental approach, which was to enable use of native patient isolates in mice. Indeed, a recent study that mouse-adapted the Spike protein was also insufficient to recapitulate severe human disease, suggesting, as in the experience with MERS-CoV, that other viral elements also need to be mouse adapted to cause COVID-19-like disease (Gu et al., 2020).

We thus hypothesized that the use of a native murine beta coronavirus may obviate many of the limitations inherent in murinizing SARS-CoV-2 and humanizing mice, better recapitulate human disease, and, importantly, be a broadly useful BSL2 platform widely accessible to the research community to explore COVID-19 pathophysiology, recognizing that it may not recapitulate some host-pathogen features specific to SARS-CoV-2. To this end, we utilized the murine coronavirus Mouse Hepatitis Virus (MHV) Strain A59. MHV-A59 is a positive-sense virus with a single-stranded RNA genome of approximately 31 kilobases which bears high similarity to SARS-CoV-2. MHV-A59 has been previously shown to cause pneumonia in C57Bl6/J mice (Coronaviridae Study Group of the International Committee on Taxonomy of, 2020; Yang et al., 2014), unlike other MHV strains that require A/J, C3H/HeJ, or other backgrounds to develop pneumonia (DeAlbuquerque et al., 2006; Leibowitz et al., 2010). As seen in Supplemental Figure 1A, SARS-CoV-2 and MHV-A59 belong to the same phylogenetic clade (adapted from (Gorbalenya et al., 2004)), have highly similar genomic structure (Supplemental Figure 1B) (adapted from (Frieman and Baric, 2008; Naqvi et al., 2020)) and share high degrees of peptide sequence homology (Supplemental Figure 1C). MHV-A59 utilizes the entry receptor CEACAM1, which is expressed on respiratory epithelium, but also on enterocytes, endothelial cells, monocytes, and neurons, as is the case with ACE2 (Compton et al., 1992; Godfraind et al., 1995). To assess the similarities in tissue tropism, we found that *Ceacam1* and *Ace2* are co-expressed in most tissues that we analyzed. While *Ceacam1* is more highly expressed in liver, *Ace2* is more highly expressed in kidney, with very similar expression in other assessed organs (Supplemental Figure 1D), and both genes were similarly regulated following respiratory MHV-A59 infection consistent with reports that both genes are interferon-inducible (Supplemental Figure 1E) (Dery et al., 2014; Ziegler et al., 2020). Therefore, we intranasally infected C57Bl6/J mice with MHV-A59 to test whether it could serve as a model of COVID-19.

**Figure 1.**
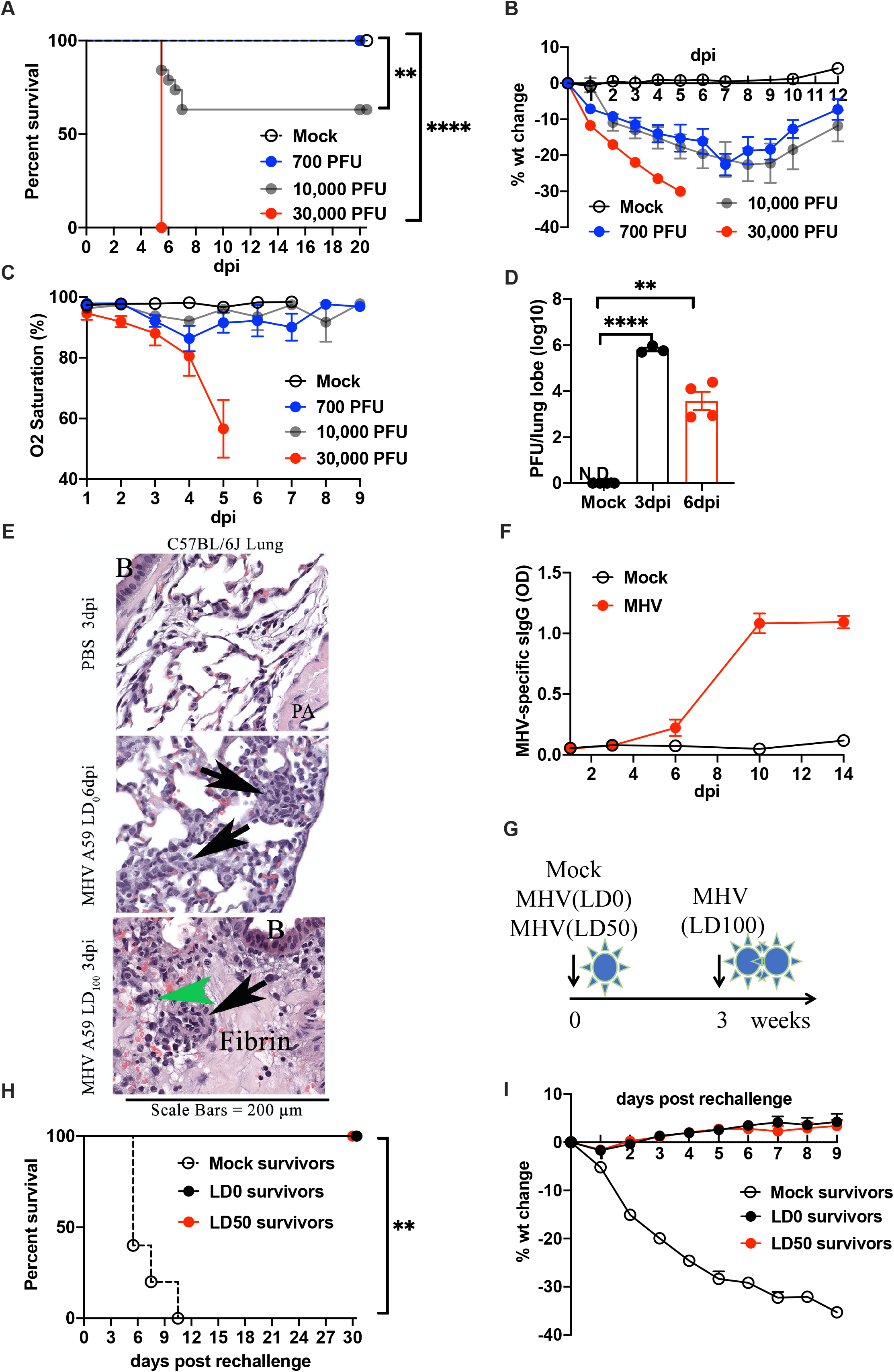
Respiratory MHV-A59 infection leads to severe clinical disease and immunological memory. **(A)** Survival rate, **(B)** Percentage of body weight change, **(C)** Oxygen saturation from C57BL6/J mice intranasally inoculated with MHV-A59 at indicated PFU. **(D)** Quantification of viral load via PFU assay from the left upper pulmonary lobe. Values were log_10_ transformed. N.D. non-detectable. **(E)** Representative images of immunohistochemistry staining of lungs from C57BL6/J mice infected with MHV-A59 at indicated doses or Mock control (PBS). Black arrows demonstrated foci of alveolar inflammation, green arrowheads indicated MHV syncytia. B = Airway Bronchi; TB = terminal bronchiole; PA = pulmonary artery. **(F)** Kinetics of serum MHV-A59-specific IgG from C57BL6/J mice infected with unlethal dose of MHV-A59 or Mock control. **(G)** Schematic of rechallenge experiment. C57BL6/J mice were rechallenged with a lethal dose three weeks post the first infection of MHV-A59 at LD_0_, LD_50_, or a Mock control. **(H)** Survival rate, **(I)** Percentage of body weight change of convalescent mice rechallenged with the lethal MHV-A59. Data represented two independent experiments, 5 mice per group for each experiment. dpi, days post infection. One-way analysis of variance (ANOVA) followed by Dunnett test was applied for multiple groups statistical calculation. Results were presented as mean ± SEM. ns, not significant, \**p*<0.05, \*\**p*<0.01, \*\*\**p*<0.001, \*\*\*\**p*<0.0001.

We found that respiratory infection of MHV-A59 caused a broad spectrum of clinical symptoms that parallels the clinical heterogeneity seen in COVID-19. At high viral loads, infection caused uniform lethality (LD_100_) secondary to acute respiratory distress syndrome (ARDS). At lethal dose 50% (LD_50_) viral loads, we observed, surprisingly, that survivors and non-survivors did not differ significantly in the degree of hypoxemia, suggesting causes of death resultant from other organ system failures. Concordantly, in these animals, we observed systemic inflammation and multi-organ involvement that mirrored pathophysiological features seen in humans. We found that tissue inflammation and damage occurred at the sites of MHV-A59 dissemination distal to the respiratory portal of entry. We thus examined the roles of type-I and type-III interferons in controlling viral dissemination. Surprisingly, we found that type I interferon receptor-deficient (IFNAR1^−/−^) animals, but not type III interferon-deficient mice, were exquisitely sensitive to MHV-A59-induced mortality but that it was not resultant from respiratory failure. Indeed, IFNAR^−/−^ animals displayed minimal hypoxemia and little enhanced pulmonary damage, viral burden, and inflammation. Instead, we found that type-I interferon was required for limiting systemic dissemination of MHV-A59, and, specifically, limiting dissemination and inflammation to other organs and in regions of the brain, including the hypothalamus, necessary for maintaining autonomic control. Consistent with these observations, early treatment with intrapulmonary recombinant type-I interferon significantly improved survival of MHV-A59-infected animals, while later treatment had no effect. Here we establish a native mouse betacoronavirus preclinical model of COVID-19 that may be useful for the study of COVID-19 pathophysiology and reveal the potential therapeutic role for type I interferons in this disease. This work contributes to the urgent need for early detection and intervention and the development of effective antiviral agents.

## Results

### Intranasal MHV-A59 infection leads to severe and dose-dependent clinical disease

SARS-CoV-2 respiratory infection in humans leads to a large diversity of clinical manifestations. To test if respiratory MHV-A59 infection recapitulated this aspect of human disease, we intranasally inoculated C57Bl/6J animals with increasing plaque-forming units (PFU) of MHV-A59. We observed a dose-dependent effect of MHV-A59 inoculation load on rates of host lethality (Figure 1A). Interestingly, regardless of the viral inocula, animals uniformly lost weight, peaking at approximately 25% body weight loss at day post-infection (dpi) 7 (Figure 1B), which was accompanied by infection-induced anorexia (Supplemental Figure S2A). To assess pulmonary function, we monitored pulse oximetry daily in infected animals and found that LD_100_ doses of MHV-A59 lead to rapid and progressive hypoxic respiratory failure (Figure 1C), which was not uniformly observed in LD_50_ animals. Indeed, LD_100_-challenged animals uniformly had oxygen saturations below 80% and were all moribund or dead by dpi 5. Considerable heterogeneity of pathology was observed in hypoxic animals (Supplemental Figure S2F). However, histopathologic analyses of lungs in hypoxic animals sacrificed at the height of viral load demonstrated pathology consistent with ARDS, including organizing pneumonia, interstitial pneumonitis, hyaline membrane formation and type-II pneumocyte hyperplasia and occasional hemorrhage and microthrombi when assessed by a blinded pathologist (CJB, Figure 1E, and Supplemental Figure S2F). These animals were also progressively unable to regulate heart rate and body temperature, suggesting severe autonomic dysfunction, while both LD_0_ and LD_50_-infected animals, on average, showed a trend towards relative bradycardia during the course of infection (Supplemental Figure S2B-E). We thus examined the pathogen burden in the lungs of animals challenged with LD_50_ doses and found that MHV-A59 proliferated in the lungs; the viral load started to significantly decrease at the inflection point of weight loss (Figure 1D). Thus, as seen in COVID-19, respiratory infection with MHV-A59 can lead to ARDS, the severity of which, in this model, occurs as a function of viral load.

In COVID-19, it is clear that patient characteristics, such as age, male sex, and those with underlying co-morbid conditions significantly increase susceptibility to both pulmonary and extrapulmonary disease (Grasselli et al., 2020; Mallapaty, 2020; Takahashi et al., 2020; Williamson et al., 2020). In the current study, we use lean 8-12-week old (adult) regular chow-fed males. As with COVID-19, females were significantly more resistant to MHV-A59 infection than males (Supplemental Figure S2G). This is the opposite of what is reported with influenza A virus infection in mice where female mice are significantly more sensitive to influenza-mediated death (Klein et al., 2012).

Next, we evaluated whether MHV-A59 animals generated an adaptive immune response that could protect from a subsequent fatal re-challenge. As expected, infected animals generated a robust IgG response to MHV-A59 (Figure 1F), and by 14 dpi had high titers of most IgG subclasses (Supplemental Figure S2H-J). Re-challenging animals that survived the initial infection 21 dpi with LD_100_ doses led to complete protection regardless of the original inoculating dose (Figure 1G, H). Impressively, convalescent animals did not lose any weight (Figure 1I) or develop anorexia (Supplemental Figure 2K). This protection was specific to MHV-A59 as it was not observed when convalescent animals were re-challenged with respiratory infection with influenza (IAV); neither prior-nor co-exposure to non-lethal doses of MHV-A59 enhanced IAV-mediated mortality (Supplemental Figure 2L-P).

Taken together, our findings demonstrate that intranasal MHV-A59 causes ARDS, hypoxic respiratory failure, and death in C57Bl6/J mice as a function of the initial viral inocula. We show that virus-specific humoral immunity is generated and capable of protecting animals from LD_100_ ARDS-mediated fatal viral loads. We also did not observe obvious synergy between IAV and MHV-A59 in enhancing mortality.

### Respiratory MHV-A59 infection causes systemic inflammation and multiorgan damage as a function of systemic viral dissemination

We were intrigued by the phenotype observed in LD_50_-challenged animals, since they did not demonstrate clear hypoxic respiratory failure, and yet half succumbed to the respiratory infection, as it was highly reminiscent of the clinical heterogeneity seen in human COVID-19. Indeed, the degree of hypoxia did not correlate with the mortality of LD_50_-challenged animals (Figure 2A), nor did it correlate with derangements in other vital signs (Supplemental Figure S3A).

**Figure 2.**
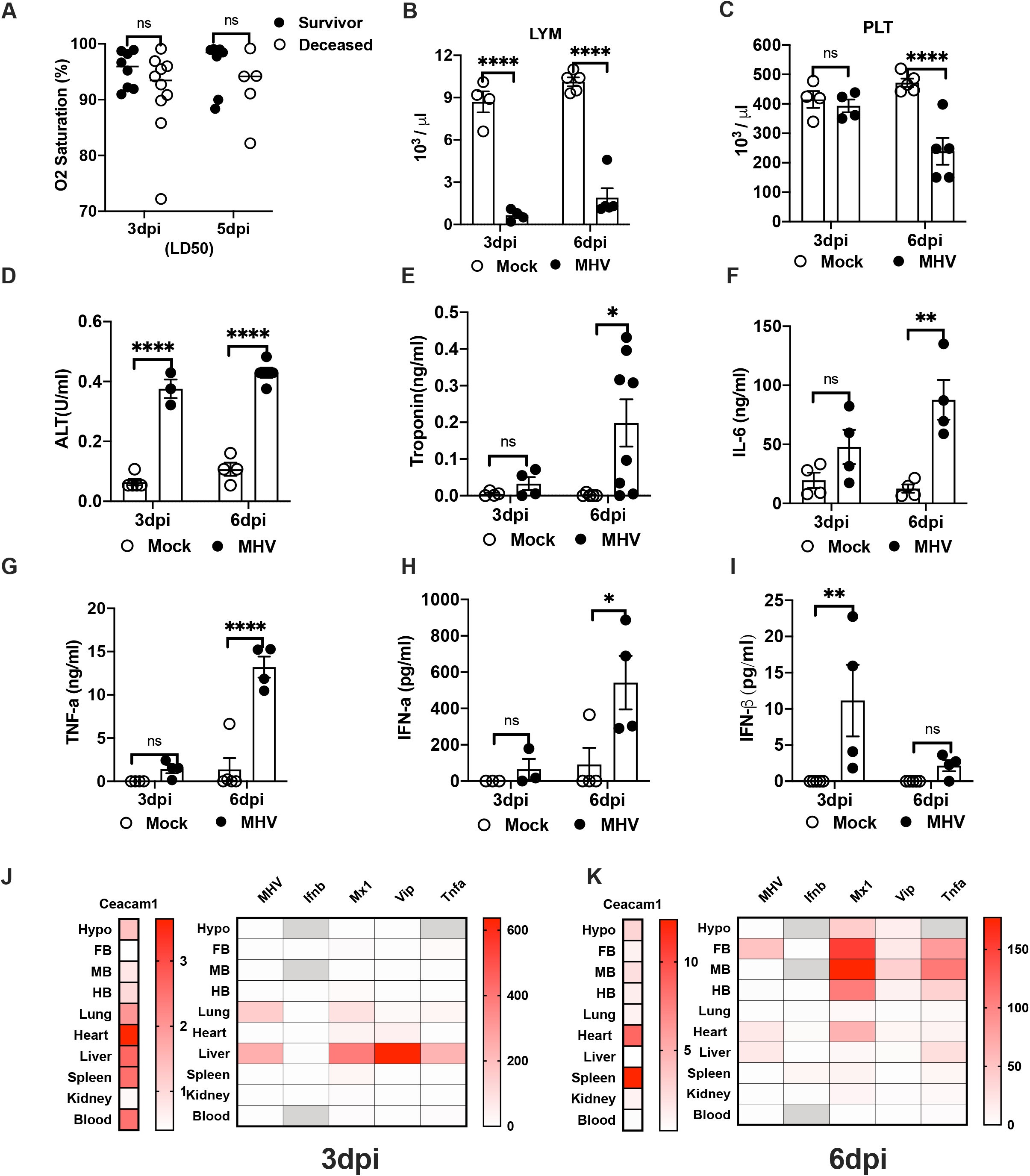
Respiratory MHV-A59 infection causes systemic inflammation and multiorgan damage associated with viral dissemination. **(A)** Oxygen saturation from surviving or deceased C57BL6/J mice on 3 or 6 days post MHV-A59 infection at LD_50_. **(B)** Quantification of lymphocytes (LYM), **(C)** platelets (PLT) in the circulation of MHV-A59 infected mice. **(D)** Serum levels of alanine transaminase (ALT), **(E)** Cardiac troponin I, **(F)** Interleukin 6 (IL-6), **(G)** Tumor necrosis factor alpha (TNF-a), **(H)** Interferon alpha (IFN-a), **(I)** Interferon beta (IFN-β) from C57BL6/J mice inoculated with MHV-A59 at LD_50_. **(J-K)** Heat map of transcriptional expressions of Ceacam1, MHV-A59, Ifnb, Mx1, Viperin (Vip), and Tnfa in tissues from LD_50_-challenged C57BL6/J mice on 3dpi and 6dpi. Viral load was indicated by relative expression of MHV-A59 compared with 18s gene given the undetectable values from mock controls. Ceacam1, Ifnb, Mx, Viperin, and Tnfa were presented as fold increase relative to mock controls. Each cell presented the mean of 8 to 12 replicates from two independent experiments. Undetectable values were shown in silver color. Hypo, hypothalamus. FB, forebrain. MB, midbrain. HB, hindbrain. Data represents two independent experiments, 4 to 6 mice per group for each experiment. dpi, days post infection. Two-way ANOVA followed by Tukey’s multiple comparisons test was applied for statistical calculation. Results are presented as mean ± SEM. ns, not significant, \**p*<0.05, \*\**p*<0.01, \*\*\**p*<0.001, \*\*\*\**p*<0.0001.

We thus examined the function of other organs in animals challenged with LD_50_ doses. We found that animals developed hematologic abnormalities (Figure 2B, C, Supplemental Figure 2Q-U) characterized by absolute lymphopenia with delayed thrombocytopenia, a feature of COVID-19 highly associated with mortality (Zhao et al., 2020). Animals also developed evidence of hepatic damage (Figure 2D), another common finding in COVID-19 associated with mortality (Hundt et al., 2020). Based on serum troponin elevation, some animals also showed evidence of cardiac injury (Figure 2E), which is a common manifestation in COVID-19 patients (Momtazmanesh et al., 2020). All animals displayed evidence of systemic inflammation (Figure 2F-I), reminiscent of a “cytokine storm,” similar to what is observed in COVID-19 (Lucas et al., 2020; Ragab et al., 2020); circulating levels of tumor necrosis factor-α (TNF-α), interleukin-6 (IL-6), and interferon-α (IFN-α) peaked at 6 dpi, when viral loads in the lung were decreasing, except for interferon-β (IFN-β), which peaked at 3 dpi.

In autopsy series, SARS-CoV-2 has been shown to variably disseminate to extrapulmonary organs where it is thought to drive extrapulmonary manifestations of COVID-19 immunopathology (Gupta et al., 2020; Puelles et al., 2020). Indeed, part of the clinical heterogeneity in COVID-19 is thought to stem from the wide expression of ACE2 on cells outside the lung and resultant viral dissemination and local injury resulting from direct viral damage or associated immunopathology. As shown, ACE2 and CEACAM1 are co-expressed in many organs (Supplemental Figure 1D-E). We found that even at LD_0_ doses, respiratory infection with MHV-A59 led to viral dissemination to multiple organs, including parts of the brain (Supplemental Figure S3B). Thus, to determine if organ damage at the LD_50_ dose correlated with viral dissemination, we assessed for inflammation and MHV-A59 burden across multiple tissues (Figure 2J, K and Supplemental Figure 3C-N). We found that infection with MHV-A59 induced expression of *Ceacam1* in many tissues (Figure 2J, K, left). On 3 dpi, there was a significant increase in viral burden in the liver and lung, which corresponded with an induction of canonical interferon-stimulated genes (ISG) and inflammatory transcripts (Figure 2J). On 6 dpi, there were significant increases in viral burden in the heart and brain, with corresponding increases in ISGs and inflammatory transcripts (Figure 2K and Supplemental Figure S3H-K). These results are consistent with plasma biomarkers indicating hepatic and cardiac damage. Interestingly, there was not a strong correlation between *Ceacam1* relative abundance (Supplemental Figure 1D) or fold induction by MHV-A59 infection, similar to what is observed in viral dissemination relative to *Ace2* expression in humans (Gupta et al., 2020; Puelles et al., 2020). In our studies, animals were systemically perfused for tissue fixation so that our findings are unlikely to reflect a significant contribution of MHV by peripheral blood to the measured content in tissues. However, we observed a strong correlation between tissues in which there was high MHV-A59 burden and tissue inflammation, which corresponded to tissues where we had observed damage, including parts of the brain required for autonomic control, the liver, and heart.

**Figure 3.**
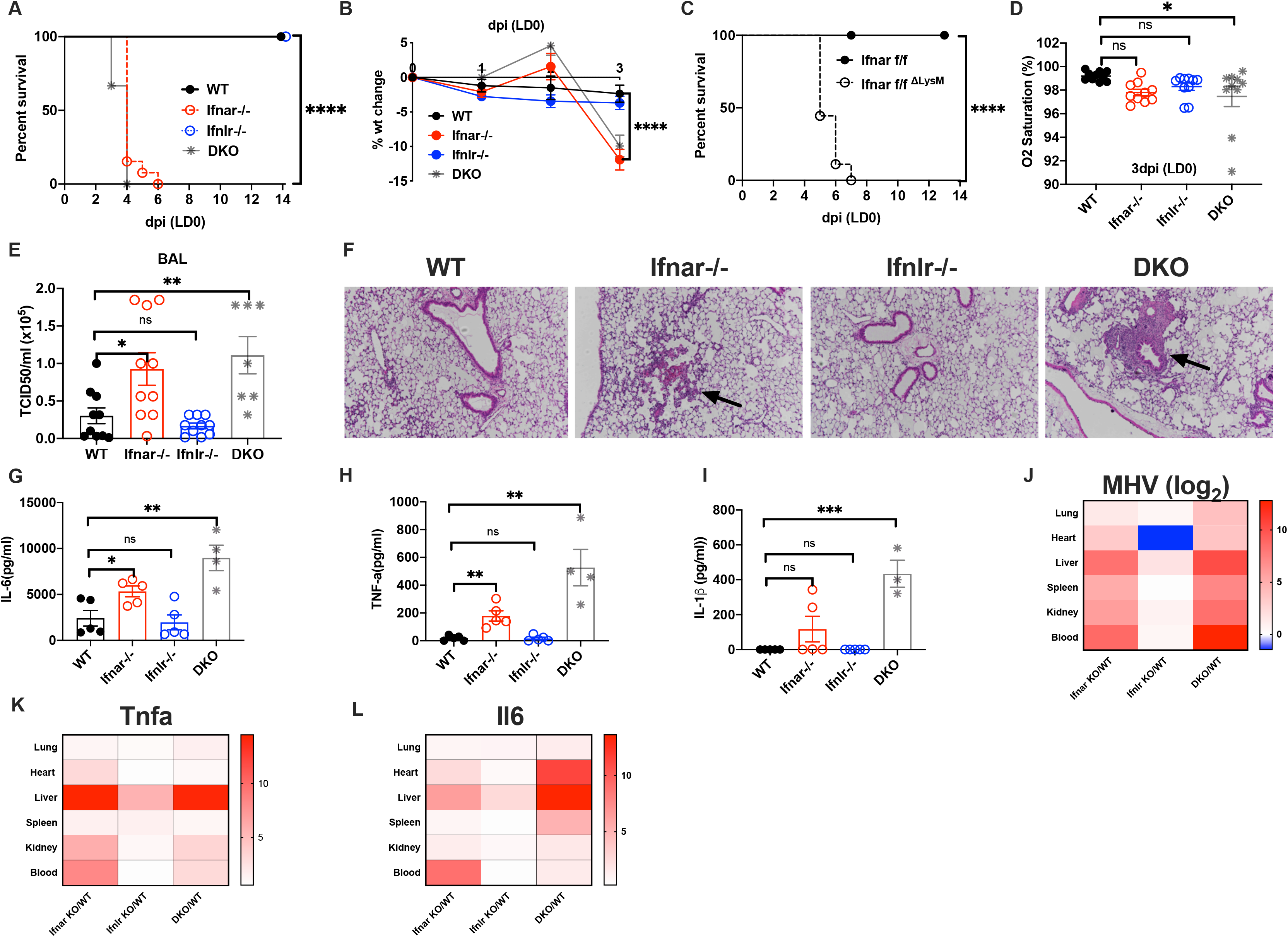
Absence of the type-I Interferon receptor during respiratory MHV-A59 infection results in viremia and mortality. **(A)** Survival rate, **(B)** Percentage of body weight change from respiratory MHV-A59 infection mice with deletion of IFNAR (Ifnar−/−), IFNLR (Ifnlr−/−), or double knockout (DKO), and wildtype (WT) controls. MHV-A59 were inoculated intranasally at an unlethal dose (LD_0_) tested from C57BL6/J mice. **(C)** Survival rate from respiratory MHV-A59 infection mice with myeloid cell specific deletion of IFNAR (Ifnar f/f ^ΔLysM^) or littermate controls (Ifnar f/f). (D) Oxygen saturation, **(E)** 50% tissue culture infectious doses (TCID50) of the virus in the bronchoalveolar lavage (BAL) from Ifnar−/−, Ifnlr−/−, DKO, or WT mice measured 3 days post inoculation of MHV-A59 at LD_0_. **(F)** Representative images of immunohistochemistry staining of lungs from indicated mice on 3dpi with LD_0_ MHV-A59 infection. **(G)** Serum levels of IL-6, **(H)** TNF-a, **(I)** IL-1β on 3dpi from indicated mice infected with LD_0_ MHV-A59. **(J)** Heat map of transcriptional expressions of MHV-A59, **(K)***Tnfa*, and **(L)***Il6* in tissues from LD_0_-challenged mice on 3dpi. Data were plotted as fold changes of expression of MHV-A59, Tnfa, and Il6 from interferon receptor knockout mice relative to WT controls. The ratio of MHV-A59 expression between genetically modified mice and WT controls were log_2_ transformed and presented in logarithmic scale. Each cell is a mean of 12-15 replicates from two independent experiments. Data represents two independent experiments, 4 to 6 mice per group for each experiment. dpi, days post infection. One-way analysis of variance (ANOVA) followed by Dunnett test was applied for multiple groups statistical calculation. Results are presented as mean ± SEM. ns, not significant, \**p*<0.05, \*\**p*<0.01, \*\*\**p*<0.001, \*\*\*\**p*<0.0001.

Our data thus suggests that extrapulmonary organ damage was resultant from viral dissemination and associated immunopathology in animals given doses of MHV-A59 insufficient to uniformly cause respiratory failure. Importantly, respiratory infection of MHV-A59 captures a surprising number of prognostic and characteristic clinical findings in COVID-19.

### Type-I Interferons are Necessary for Pathogen Control in Respiratory MHV-A59 Infection

Type-I and III interferon (IFN)-mediated responses play important roles in the viral immune response, both in controlling pathogen burden and also in tuning the immune response (Grajales-Reyes, 2020; Park and Iwasaki, 2020). Decreased Type-I IFN response has been correlated with increased disease severity in human SARS-CoV-2 patients (Hadjadj et al., 2020), leading to clinical trials of recombinant IFN-β in COVID-19 patients (Walz et al., 2020). Given that we observed variable viral dissemination in LD_50_-challenged animals as a pathogenic cause of mortality, we thus examined the role of Type-I and III interferons in the host response to MHV-A59.

We inoculated mice deficient in either type-I interferon receptor (IFNAR1^−/−^) or type-III interferon receptor (IFNLR1^−/−^), or mice deficient in both of these receptors (DKO). Indeed, one major advantage of the MHV-A59 model is that it causes COVID-19-like disease on a C57Bl56/J genetic background, allowing for the use of readily available genetic tools for pathway dissection. We found that IFNAR1^−/−^ and DKO mice were exquisitely sensitive to MHV-A59. LD_0_ PFUs in wildtype mice caused 100% lethality in IFNAR1^−/−^ and DKO animals, which succumbed to infection by 4 dpi (Figure 3A), regardless of gender of animal (Supplemental Figure S4A). This was associated with a modest change in weight loss significant at 3 dpi (Figure 3B). No apparent morbidity or mortality was observed in IFNLR1^−/−^ animals, suggesting a major role for IFNAR1 and a dispensable role of IFNLR1 in mediating host survival to MHV-A59 infection at this dose. However, using a viral inoculum of 100 PFU (100-fold dilution of LD_0_ for wildtype animals), we observed increased mortality in the DKO compared to IFNAR1^−/−^ mice, unveiling a potential protective role of type-III interferon (Supplemental Figure S4B).

Since IFNAR is expressed in virtually all cells in the body, we next asked if IFNAR1 expression would be required in the myeloid compartment, as had been previously demonstrated when MHV-A59 was inoculated via the intraperitoneal route (Cervantes-Barragan et al., 2009). We thus generated mice with myeloid-specific deletion of IFNAR1 using Cre-mediated deletion under the control of the Lysozyme 2 promoter (LysMCre, Ifnar f/f ^ΔLysM^). As expected, Ifnar f/f ^ΔLysM^ animals were significantly more susceptible to MHV-A59 infection than their f/f littermate controls, indicating that IFNAR expression was required on myeloid cells (Figure 3C).

We then examined respiratory function in type-I and type-III interferon-deficient animals. Unexpectedly, IFNAR1^−/−^ and DKO animals did not display significant hypoxemia when moribund (Figure 3D) despite modestly increased viral burdens and pleocytosis in the bronchoalveolar lavage (BAL) (Figure 3E and Supplemental Figure S4C). This was consistent with relatively minor observed histopathologic evidence of pulmonary disease (Figure 3F). Next, we looked at the magnitude of systemic inflammation. We observed modest increases in systemic inflammatory cytokines at the same peri-moribund time point (Figure 3G-I). We did not detect differences in the amount of MHV-A59-specific IgM levels as a result of IFNAR1 deficiency (Supplemental Figure S4D).

However, when we quantitated viral burden in a wide range of organs, we found that MHV-A59 burden was dramatically increased in virtually every tissue we assessed in animals lacking IFNAR1, including the blood (Figure 3J log-scale, and Supplemental Figure S4E-K, linear-scale), which corresponded with increased inflammatory transcripts (Figure 3K-L and Supplemental Figure S4L-S, linear-scale). While all organs from IFNAR1^−/−^, but not IFNLR1^−/−^, animals displayed >10 fold more viral load compared to wildtype controls, the heart and liver displayed the most increase in inflammatory transcripts, with greater than 10-fold increases in the observed amount of *Il6* message compared to wildtype.

Taken together, MHV-A59 infection results in induction of type-I interferons which is required for controlling systemic dissemination of pathogen burden. LD_0_ doses are uniformly and rapidly lethal in IFNAR1^−/−^ animals. Unexpectedly, IFNAR1 deficiency does not significantly lead to enhanced pulmonary viral burden or death resultant from hypoxemic respiratory failure.

### MHV-A59 respiratory infection in IFNAR-deficiency reveals vulnerability in autonomic control

To assess the cause of death in IFNAR-deficient animals, we assessed organ function and vital signs. We focused on the liver and heart given the dramatic changes in inflammatory transcripts observed in these organs. Unexpectedly, despite evidence of increased hepatic and cardiac viral load, we observed neither significantly increased biomarkers (ALT and troponin) indicative of organ injury (Figure 4A and B, Supplemental Figure S5A), nor histopathological changes consistent with significant increased tissue damage in IFNAR1^−/−^ versus wildtype controls (Supplemental Figure S5B, C). Consistent with the correlation between lymphopenia and worse outcomes, IFNAR1^−/−^ animals displayed more lymphopenia (Supplemental Figure S5D) than wild-type controls

**Figure 4.**
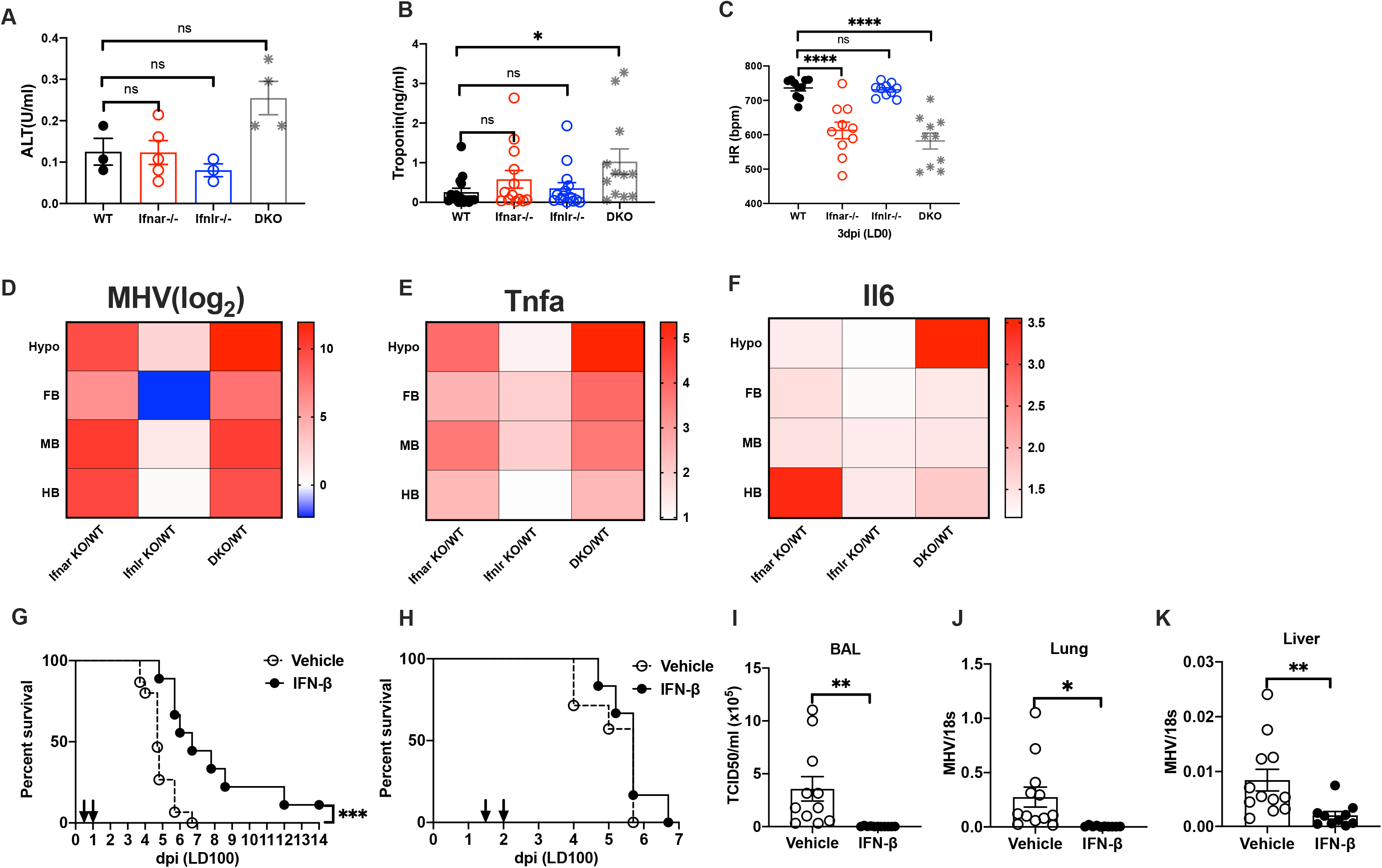
Type-I interferon receptor deficiency reveals vulnerability to CNS metastasis and failure of autonomic control. **(A)** Serum levels of alanine transaminase (ALT), **(B)** Cardiac troponin I, and **(C)** Heart rate (HR) from genetical modified or wildtype mice on 3 dpi with intranasal inoculation of MHV-A59 at LD_0_. **(D)** Heat map of transcriptional expressions of MHV-A59, **(E)** Tnfa, and **(F)** Il6 in tissues from LD_0_-challenged mice on 3dpi. Data were plotted as fold changes of relative expression of MHV-A59, Tnfa, and Il6 from interferon receptor knockout mice with respect to that from WT controls. The ratio of MHV-A59 expression between genetically modified mice and WT were log_2_ transformed and presented in logarithmic scale. Each cell presented mean of 10 replicates from two independent experiments. Hypo, hypothalamus. FB, forebrain. MB, midbrain. HB, hindbrain. **(G-H)** Survival rate of C57BL6/J mice intranasally inoculated with lethal MHV-A59 at LD_100_ followed by **(G)** early or **(H)** late treatment of interferon-β (IFN-β). IFN-β was applied at 12 and 24 hours post infection for early phase, or at 24 and 36 hours post infection for late phase treatment. Arrows indicated the timepoints for IFN-β intratracheal application. **(I)** 50% tissue culture infectious doses (TCID50) of the virus in the bronchoalveoar lavage (BAL), relative expression of MHV-A59 in **(J)** the lung and **(K)** the liver from MHV-A59 infected mice received early IFN-β treatment as indicated in (G). Data represent two independent experiments, 5 to 6 mice per group for each experiment. One-way analysis of variance (ANOVA) followed by Dunnett test for multiple groups, or unpaired t-test for two groups was applied respectively for statistical calculation. Results are presented as mean ± SEM. ns, not significant, \**p*<0.05, \*\**p*<0.01, \*\*\**p*<0.001, \*\*\*\**p*<0.0001.

However, we detected diminished heart rate responses to viral infection in IFNAR-deficient animals (Figure 4C) but not in other vital parameters, including pulse distension, which is a surrogate for blood pressure (Supplemental Figure S5E-H). Given the high infectivity of MHV amongst mouse colonies, it was not possible in our animal facilities to perform continuous invasive telemetric monitoring of blood pressure and heart rate to better assess heart rate variability or blood pressure. Maintenance of heart rate is in-part controlled by central nervous system (CNS), and especially the hypothalamus and brainstem (Farmer et al., 2016; Pinol et al., 2018). We thus assessed MHV-A59 burden in different parts of the brain and observed profound increases in viral load in the brain, including the hypothalamus and brainstem in IFNAR1-deficient mice relative to wildtype controls (Figure 4D, Supplemental Figure S4E). Analyses of single-cell RNA sequencing of the hypothalamus demonstrates nearly identical co-expression of *Ace2* and *Ceacam1* (Supplemental Figure S5I). This was associated with corresponding increases in tissue inflammatory transcripts (Figure 4E-F). These findings are consistent with our findings that LD_50_ animals displayed central nervous system inflammation, including CNS inflammation (Figure 2K), and relative bradycardia generally displayed by infected animals, regardless of inoculating dose, throughout their course of disease (Figure S2C). Our data thus suggests that MHV-A59-induced CNS dysfunction may be a key component in pathogenesis, a vulnerability that was magnified and revealed in the context of IFNAR deficiency. Our findings in the CNS are reminiscent of the ‘long-haulers’ phenotype, where many convalescent COVID-19 patients continue to have symptoms of dysregulated autonomic control (Romero-Sanchez et al., 2020).

Having established a role of interferon deficiency as a major contributor to viral dissemination and subsequent extrapulmonary damage and resultant death, we aimed to understand if type-I interferon treatment could improve viral clearance in MHVA-59 infection. This therapeutic is currently being pursued in clinical trials (Walz et al., 2020). To test this, we treated mice with interferon β at 1μg/mouse by intra-tracheal route at early (12 hours and 36 hours) or later (36 hours and 48 hours) timepoints after a lethal dose of MHV-A59 intranasal infection. Mice that were treated early but not later in the disease course were significantly protected from MHV-A59 infection (Figure 4G, H). As expected, early treatment of IFN-β suppressed viral production as indicated by the decreased viral load in the BAL and lung tissue (Figure 4I, J) and in the liver (Figure 4K) from infected animals 3 days post infection. Consistent with our knockout studies, protection was also not observed with IFN-λ administration at early disease course (Supplemental Figure S5J). Finally, we performed convalescent serum transfers in wildtype and IFNAR1^−/−^ animals and found that a single low-volume injection of convalescent serum provided modest but significant protection from MHV-A59 induced death and reduced viral load (Supplemental Figure S5K-O), suggesting adaptive immunity can provide some protection despite major deficiency in innate immune system.

Taken together, we demonstrate that respiratory infection with a native mouse betacoronavirus, MHV-A59, recapitulates most aspects of COVID-19 pathophysiology and clinical heterogeneity. We demonstrate its utility as a BSL2 platform on the C57/Bl6J genetic background to dissect disease pathogenesis and as a useful pre-clinical model for testing COVID-19 therapeutics. This work also demonstrates the utility of type I interferon as a potential therapeutic option against coronaviral infections, which are critically needed in COVID-19.

## Discussion

As the COVID-19 pandemic continues to spread across the globe, the need to understand disease pathophysiology and to apply this knowledge to novel therapeutic strategies is an urgent need. Animal models are critical in this effort, with mice being the most efficient models given their size, ease of husbandry and maintenance, as well as the vast available genetic and pharmacologic tools. In the current study, we show that C57Bl6/J mice infected with a native murine betacoronavirus MHV-A59 recapitulates many aspects of COVID-19 pathophysiology. Intranasal inoculation of MHV-A59 produced a productive infection that develop many of the hallmark features of human COVID-19 disease, including ARDS, lymphopenia, multiorgan involvement, and systemic inflammation. We thus present a BSL2 platform animal model using C57Bl6/J that may be a useful tool in the ongoing COVID-19 effort. This model could be widely available to laboratories given a lack of need for BSL3 facilities and the use of the most commonly used inbred strain and greatly enhance COVID-19-related efforts.

Our model utilizes MHV-A59, a beta coronavirus highly phylogenetically related to SARS-CoV-2 that is native to mice, and the scourge of mouse colonies given its high infectivity to case fatality rate (Compton et al., 2004). The major limitations of MHV-A59 is that it 1) does not capture all SARS-CoV-2-intrinsic pathogenic factors and that 2) the entry receptor of MHV-A59 CEACAM1 is not co-localized exactly with that of ACE2-expressing cells. However, we argue that, teleologically, our approach may not be that disparate from the ongoing efforts attempting to humanize mice and murinize SARS-CoV-2 to recapitulate human disease. Indeed, the only models that currently recapitulate severe disease using SARS-CoV-2 require the forced expression (using K18 or Hfh4 promoters) of human ACE2 in tissues which do not endogenously express ACE2 (Jiang et al., 2020; McCray et al., 2007). Likewise, based on the published data and previous experience with MERS-CoV, it is highly likely that murinization of SARS-CoV-2 will be required in order to produce COVID-19-like disease in mice where endogenous ACE2 expression is maintained. Therefore, while our model may lack certain features important for SARS-CoV-2-specific host-pathogen interactions, it is unlikely that any host other than a human will capture SARS-CoV-2-specific host-pathogen interactions. On the other hand, the MHV-A59 mouse model, which is the species-appropriate pathogen in its corresponding host, recapitulates a surprising number of characteristics of COVID-19.

Perhaps the most astonishing feature COVID-19 is the wide breadth of clinical manifestations. A confounding factor in these clinical epidemiologic studies is that the time and dose of viral inoculation is unknown and thus different arrays of manifestation may simply represent differences in the inoculum size and duration of the disease course. Despite this caveat, the observed disease trajectories are widely variable and range from asymptomatic disease to death with an unpredictable array of clinical manifestations. The host substrate and viral inocula are two factors that are critical in determining the disease trajectory. The MHV-A59 model recapitulates the enhanced susceptibility of male animals to lethal disease observed in COVID-19 (Grasselli et al., 2020; Mallapaty, 2020; Takahashi et al., 2020; Williamson et al., 2020). It remains to be seen if mouse models of aging, obesity, myocardial infarction or hypertension will also be more sensitive to MHV-A59 respiratory infection and what vulnerabilities are conferred by these underlying disease states.

In the young adult human population without known underlying comorbidities, mortality seems to be stochastic (DeBiasi et al., 2020). We speculate that respiratory viral load may be a critical determinant of disease severity, in line with the observations made amongst physicians caring for COVID-19, wherein those with greatest droplet exposure either through endotracheal intubation (respiratory therapists, anesthesiologists, and emergency room doctors) or otherwise (dentists, ENT and ophthalmologists) had the highest rates of mortality (Ing et al., 2020; Nguyen et al., 2020). Indeed, as reported in the current SARS-CoV-2 mouse models, we find that the level of viral load is a key determinant of disease in the MHV-A59 model.

In our model, the trajectory of disease in mice receiving high respiratory inoculum is respiratory failure. However, in lower doses of viral inoculum this was no longer observed. Indeed, using an LD_50_ dose of MHV-A59, we found that death did not correlate with hypoxemia. Rather, disease trajectory seemed to be determined by viral dissemination and extrapulmonary multi-organ inflammation and damage. It is now well-recognized that SARS-CoV-2 is widely detectable in multiple tissues; our findings are in-line with recently reported human autopsy series of COVID-19 patients, where there is variable expression of virus in lung, heart, liver, brain, and blood (Puelles et al., 2020). Interestingly, the pattern of viral dissemination did not correlate tightly with entry receptor expression, as is also observed with ACE2 and SARS-CoV-2. CEACAM1, like ACE2, is also expressed on endothelial cells and monocytes (Ergun et al., 2000; Hamming et al., 2004), and it is likely that bulk-level detection of these entry receptors, as was done in this study, lacks the necessary resolution to capture this aspect of cellular infectivity which may underly the mechanism of dissemination. Nonetheless, our studies using LD_50_ doses indicate that despite relatively preserved lung function, extrapulmonary inflammation mediated by viral dissemination to extrapulmonary tissues may represent a common pathophysiological mechanism of action that accounts for the wide heterogeneity in disease trajectories observed in respiratory SARS-CoV-2 infections in humans and respiratory MHV-A59 infections in mice. In line with the importance of viral control, and consistent with all SARS-CoV-2 animal models to date, MHV-A59 infection also generates long-lasting and serum-transferable protective immunity to subsequent LD_100_ challenges.

From this perspective, it is likely that the degree of type-I interferon response, which could vary as a function of genetics and other host characteristics, is another major determinant in disease trajectories in SARS-CoV-2. Since IFNAR signaling primes and amplifies inflammatory responses, it is both involved in immunopathologic responses as was observed in the SARS-CoV-2 model employed by the Iwasaki group (Israelow et al., 2020) as well as viral control, as was observed in the model used by the Perlman group (Hassan et al., 2020; Sun et al., 2020a). Decreased Type-I IFN response has been correlated with increased disease severity in human SARS-CoV-2 patients (Hadjadj et al., 2020). We found that IFNAR deficiency led to profound viral dissemination. In this model, type-I interferon was absolutely required for extrapulmonary viral control. In the MHV-A59 model, despite high viral loads in certain organs, like the liver and heart, expected commensurate damage and inflammation was not detected using the method we employed in this study. It is possible that this may reflect the dual roles of IFNAR signaling, where it prevents viral propagation but can come at the expense of immunopathology. Conversely, impaired IFNAR1 signaling may reduce immunopathology at the expense of viral burden. Interestingly, IFNAR1-deficient animals did not succumb to hypoxic respiratory failure but rather to an inability to maintain autonomic control in the setting of pronounced CNS viral load and inflammation.

Neurologic involvement in COVID-19 is becoming an increasingly recognized manifestation. More than 36 percent of COVID-19 patients displayed neurologic manifestations (Montalvan et al., 2020). These include hyposmia as a characteristic and dominant feature but also include frank encephalitis, including involvement of the cardiorespiratory centers, which may bely the autonomic dysfunction observed in patients. In our study, we find that MHV-A59 can be readily detected in various parts of the brain following respiratory inoculation, including areas known to be critical for cardiorespiratory control, despite poor expression of the entry receptor, much like what is observed in SARS-CoV. We find evidence of autonomic dysregulation at all doses tested of MHV-A59 led to an impaired heart rate response to infection, and this exacerbated in the context of IFNAR-deficiency, and may have contributed to death in these animals. We cannot exclude a component of adrenal insufficiency to the impaired HR response, or the development of cardiac contractile dysfunction related to cytokine activation or endothelial injury with impaired myocardial perfusion. The mechanisms underlying neuronal invasion is a long-standing mystery in the study of neurotropic betacoronaviruses (Miura et al., 2008; Perlman et al., 1989; Phillips and Weiss, 2011) and the immune and tissue responses in these privileged tissues remain to be elucidated.

Finally, we demonstrate that early control of viral replication with interferon β or convalescent serum significantly improves MHV-A59-induced mortality, while no benefit was shown with treatment at a later course. While we did not demonstrate a clear role for type-III interferon in our current study, DKO animals demonstrated relatively more severe disease. Overall, the role of type-I interferon was dominant in these studies, and it remains to be seen if treatment with type-III interferon would play a protective role when using i.e. LD_50_ doses. Our findings resonate with the general experience in human trials where the intervention timepoint within the disease trajectory is a critical determinant of the therapeutic response. Thus, in the absence of biomarkers that can specify when a patient was infected or where they lie along their disease course, it will be challenging to design immunomodulatory or antiviral therapeutics in COVID-19, unless clinical targets emerge that are overwhelmingly protective regardless of the patient’s infection duration or their particular clinical constellation. From this perspective and based on the growing appreciation for viral dissemination to be a major pathogenic driver in many COVID-19 disease endophenotypes, rapid and concerted development of antiviral therapeutics represents an urgent and relatively unmet need.

We describe a BSL2 model for studying COVID-19 pathophysiology using mouse betacoronavirus MHV-A59. Our model captures the diverse disease trajectories seen in COVID-19 ranging from relatively asymptomatic to multiorgan involvement to respiratory failure as a function of inoculating dose and host characteristics (Figure 5). Further expedited academic and biopharmaceutical research and development on COVID-19 disease pathogenesis and pathophysiology are critical to the ongoing global therapeutic effort. The BSL2 platform presented in this report may be a useful addition to the current armamentarium of research tools.

**Figure 5.**
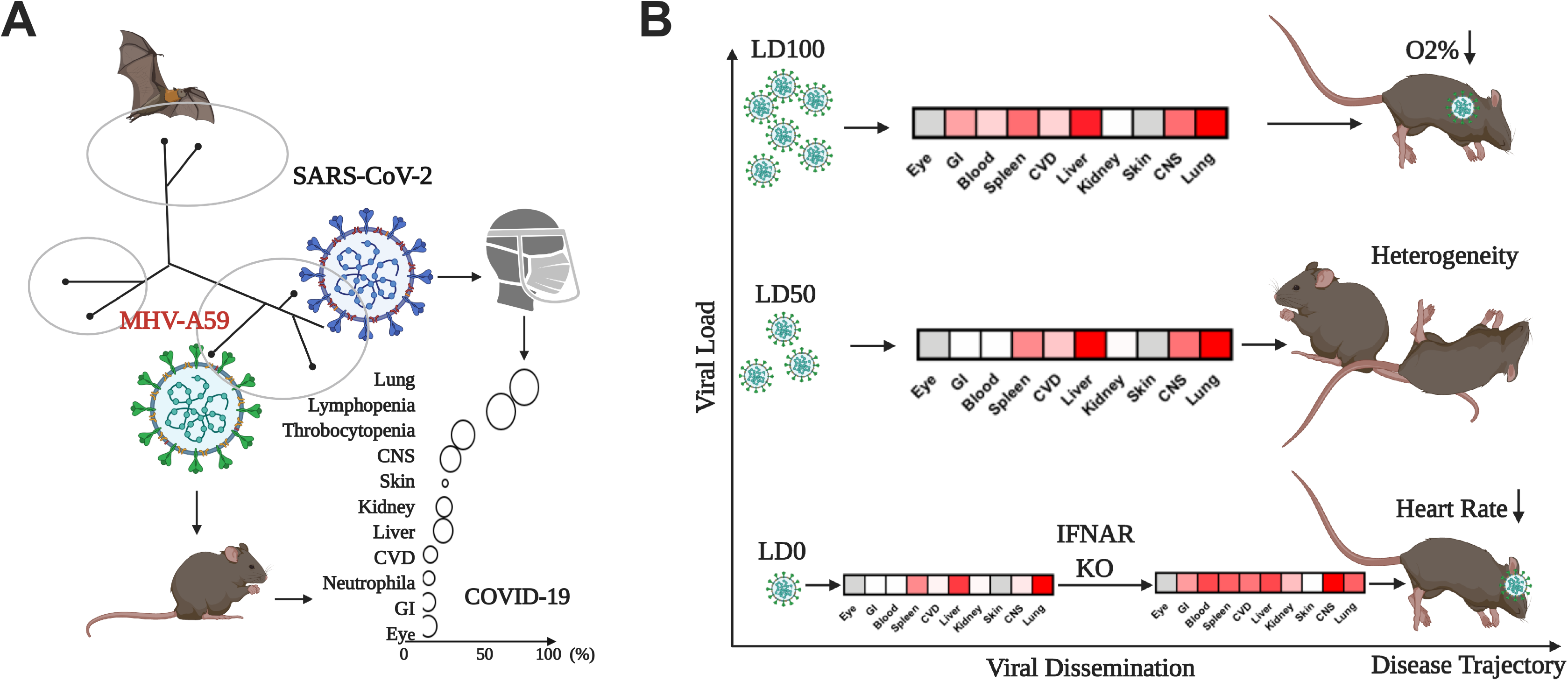
Respiratory MHV-A59 infection captures the diverse disease trajectories of COVID-19. **(A)** Homology of MHV-A59 and SARS-CoV-2 that presents organotropism beyond the respiratory system and causes multi-organ dysfunction in COVID-19 patients. The x-axis indicates the incidence of organ injury denoted in the y-axis in COVID positive patients, and the size of circles correlates with the prevalence in severe COVID-19 patients. **(B)** Illustration of the effect of viral load and host defense on different disease trajectories. High viral inoculum leads to death from ARDS and hypoxemic respiratory failure. At LD_0_ and LD_50_ inoculums, patterns of organ involvement emerge, likely as a function of viral metastasis. In the context of IFNAR-deficiency, vulnerability in the brain for viral metastasis and its consequences on autonomic control are revealed.

### Limitations of Study

The MHV-A59 model will not capture SARS-CoV-2-intrinsic host-pathogen interactions that may be pathogenic in COVID-19. Despite viral homology and entry-receptor similarities and the pathophysiologic similarities between MHV-A59-induced disease and COVID-19, this model will not capture all aspects of COVID-19. These experiments were performed in a single facility (albeit in several different animal rooms in several different laboratories within Yale), and on the C57Bl6/J background. Thus, the impact of microbiota and other facility-specific features are unknown. MHV-A59 preparations were done in the Wilen, Wang, and Compton laboratories and batch-to-batch variability will likely occur due to mutations and genetic drifts during viral propagation. The translatability of this study to humans is to be determined.

## Acknowledgments

We thank the Wang, Dela Cruz, Dixit, Pereira, Ring, and Wilen Labs for valuable discussions. We thank Yawen Jiang for technical assistance. We thank the Immunobiology and Internal Medicine Departments for solidarity in the Yale COVID-19 research effort. We thank Drs. Ruslan Medzhitov and Akiko Iwasaki for critical review of the manuscript.

The Wang lab is supported in part by NIH grant 1K08AI128745 and by generous gifts from the Pew Charitable Trusts, Knights of Columbus, G. Harold and Leila Y. Mathers Charitable Foundation, and the Ludwig Family Foundation. The Dela Cruz lab is in part supported by NIH grant HL126094, VA BX004661, DOD and Parker B. Francis award. The Dixit lab is supported in part by NIH grants P01AG051459, AR070811 and Cure Alzheimers Fund. The Pereira lab is supported in part by NIH grant AI113040. Lokesh Sharma is supported by a Parker B Francis Fellowship. The Wilen lab is supported by NIH grant K08AI128043, R01AI48467, Burroughs Wellcome Fund, Fast Grants, the Ludwig Family Foundation, and G. Harold and Leila Y. Mathers Charitable Foundation. The Young lab is supported in part by NIH grants HL150449 and HL 148344. Anush Swaminathan is supported by NIH Medical Scientist Training Program Training Grant T32GM136651.

## Author Contributions

AW, HQ, CDC, and LS conceived the project and wrote the manuscript with assistance from AS, JT, and KIW. BH propagated MHV-A59 with input from SC and CW, and also performed experiments with HQ. HQ and BH performed experiments establishing and characterizing the model system. AR contributed to the MHV-A59 model generation. LS and HQ performed experiments related to IFNAR knockouts with assistance from XP, LW and VH. SR in the laboratory of VD assisted in experiments. SK provided IFNLR−/− mice and guidance for the study. JP generated experiments related to gender differences. CZ and XP maintain animals for the Wang and Dela Cruz Labs, respectively. DL generated the ssRNASeq analyses of hypothalamus. CJB, a mouse pathologist, generated the images presented in Figure 1 and Supplemental Figure S2 and interpreted them; all other histolopathology figures and interpretations presented were done by others. XT and SI assisted in interpreting renal histology. YM and LY assisted in performing and interpreting cardiac histology.

## Supplementary Figures

**Figure S1 (Related to Figure 1).**
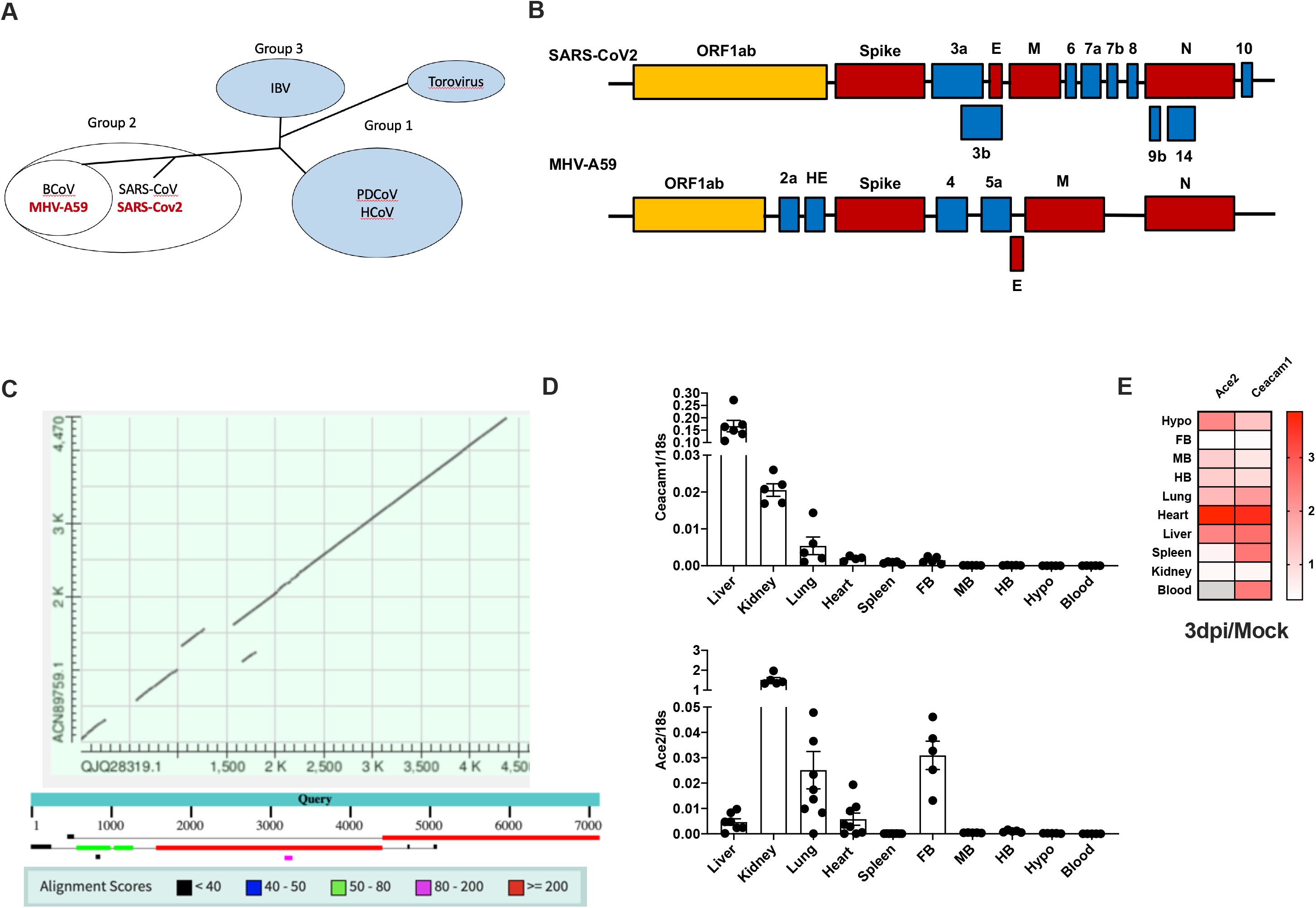
Homology between MHV-A59 and hSARS-CoV-2. Related to Figure 1. **(A)** Phylogenetic clade (IBV (avian coronavirus); BCoV (bovine coronavirus); HCoV (human coronavirus); PDCoV (porcine coronavirus)) and **(B)** genomic structures of human SARS-CoV-2 and MHV-A59. **(C)** Peptide sequence homology between human SARS-CoV-2 and MHV-A59. **(D)** Transcriptional expression of *Ace2* and *Ceacam1* in tissues from normal C57BL6/J mice. **(E)** Heat map of relative expression of *Ace2* and *Ceacam1* in tissues from C57BL6/J mice with intranasal MHV-A59 inoculation on 3 dpi. Ratio of gene expression between that from infected mice on 3 dpi and relative to that from mock controls. Each cell presented is a mean of 5 to 8 replicates from two independent experiments. Undetectable values are depicted as gray cells.

**Figure S2 (Related to Figure 1).**
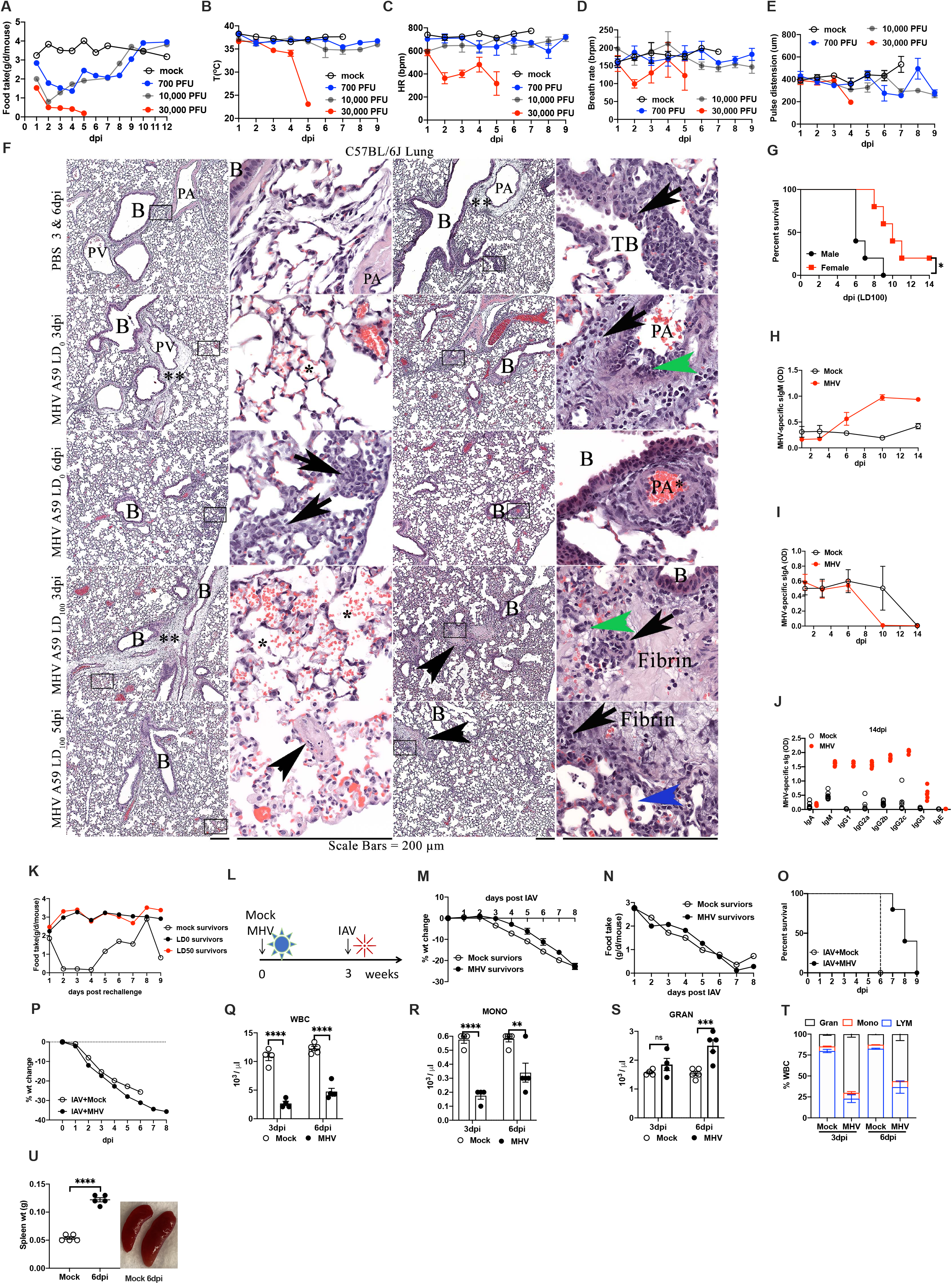
MHV-A59 infection leads to severe clinical disease and immunological memory specific to the same species, Related to Figure 1. **(A)** Food intake, **(B)** Core body temperature, **(C)** Heart rate, **(D)** Breath rate, and **(E)** Pulse distension from C57BL6/J mice intranasally inoculated with MHV-A59 at indicated PFU. **(F)** Representative HE (low power, higher power inset) from multiple C57Bl/6J male mice inoculated with intranasal (IN) PBS or MHVA59 (LD_0_ or LD_100_) that were necropsied at 3, 5, or 6 days post inoculation (dpi). Mock infected mice largely lacked pathologic findings except for mild bronchiolitis with mild perivascular/peribronchial(iolar) PV/PB edema (**) with a mild inflammatory (arrow) response, most consistent with aspiration, since the same animals also lacked pathologic changes in the liver or brain. In contrast, mice given MHV-A59 vary from none to varying degrees of PV/PB edema (**) at all doses and dpi. Alveoli vary in the presence and extent of hemorrhage (*) with more severe alveolar hemorrhage observed mice given lethal doses of MHVA59. Regardless of dose or dpi, MHVA59 mice demonstrate foci of alveolar inflammation (black arrows) with neutrophils within alveoli (blue arrowheads). Mice given a LD_100_ dose have multiple foci of fibrin (black arrowheads), hemorrhage and edema, with varying severity of lung lesions. MHV syncytia (green arrowheads) are observed as well as edema (*). * = alveolar hemorrhage; **= perivascular and/or peribronchial(iolar) edema; B = Airway Bronchi; TB = terminal bronchiole; PA = pulmonary artery; PV - pulmonary vein; arrows - alveolar hemorrhage; blue arrowhead - neutrophils in alveoli; green arrowheads - MHV Syncytial cells; black arrowheads - fibrin, hemorrhage, edema. **(G)** Survival rate of males comparing with females of C57BL6/J mice. **(H)** Kinetics of serum MHV-A59-specific IgM, **(I)** IgA from C57BL6/J mice infected with unlethal dose of MHV-A59 or Mock control. **(J)** Serum levels of indicated immunoglobulins on 14 dpi from C57BL6/J mice infected with unlethal dose of MHV-A59 or Mock control. **(K)** Food intake of survivors from MHV-A59 infection at LD_0_, LD_50_, or the mock control when they were rechallenged with a lethal dose of MHV-A59 three weeks later. **(L)** Schematic of survivors from MHV-A59 (MHV) infection challenged with influenza A virus (IAV) 3 weeks later. **(M)** Percentage of body weight change, **(N)** food intake of survivors from MHV-A59 infection challenged with IAV as indicated in (L). **(O)** Survival rate, **(P)** Percentage of body weight change from C57BL6/J mice intranasally inoculated with lethal IAV, combined with an unlethal MHV-A59 or a mock control. **(Q)** White blood cells (WBC), **(R)** Monocytes (MONO), **(S)** Granulocytes (GRAN), and **(T)** Percentage of WBC subtypes from C57BL6/J mice infected with MHV-A59 at indicated timepoint. **(U)** Splenomegaly (spleen weight, and representative picture) at 6 dpi from intranasally MHV-A59 infected C57BL6/J mice. Five mice per group for every experiment. Two-way ANOVA followed by Tukey’s multiple comparisons test, or unpaired t-test for two groups was applied respectively for statistical calculation. Results were presented as mean ± SEM. dpi, days post infection.

**Figure S3 (Related to Figure 2).**
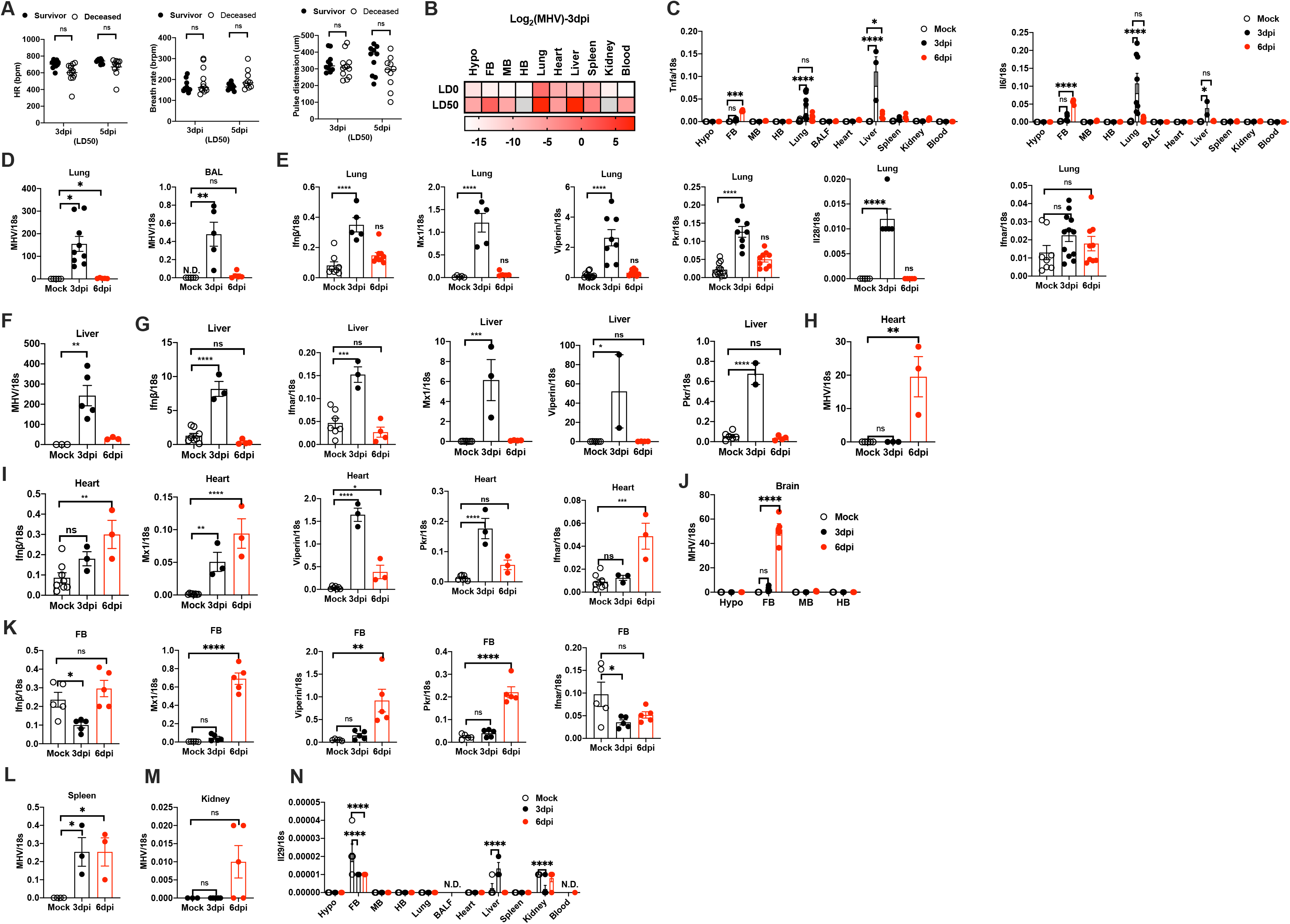
Respiratory MHV-A59 inoculation causes systemic inflammation and multiorgan damages associated with viral dissemination, Related to Figure 2. **(A)** Heart rate (HR), breath rate, and pulse distension from survivors or deceased C57BL6/J mice on 3 or 5 days post intranasal inoculation of MHV-A59 at LD_50_. **(B)** Heat map of transcriptional expressions of MHV-A59 in tissues from C57BL6/J mice 3 days post intranasally inoculated with MHV-A59 at LD_0_ or LD_50_. Viral load was presented as relative expression of MHV-A59 with log2 transformation. Each cell presented mean of 8 to 10 replicates from two independent experiments. Undetectable values were shown in silver color. Transcriptional expression of **(C)** Tnfa, and Il6 in tissues from C57BL6/J mice with intranasal MHV-A59 infection at LD_50_ at indicated time. Transcriptional expression of MHV-A59, type I interferon (Ifnβ), interferon stimulated genes (Mx1, Viperin, Pkr), type III interferon (Il28, Il29), Ifnar **in (D-E)** the lung and bronchoalveolar lavage (BAL), **(F-G)** liver, **(H-I)** heart, **(J-K)** four regions of brain, **(L)** spleen, **(M)** kidney from C57BL6/J mice with intranasal MHV-A59 infection at LD_50_ at indicated time. Hypo, hypothalamus. FB, forebrain. MB, midbrain. HB, hindbrain. dpi, days post infection. Date represented two independent experiment, 5 mice per group for every experiment. Two-way ANOVA followed by Tukey’s multiple comparisons test for statistical calculation of panel (A). One-way analysis of variance (ANOVA) followed by Dunnett test for multiple groups comparation of panel (C – N). Groups in (C, J and N) without labeling indicates no significant difference. Results were presented as mean ± SEM. ns, not significant, \**p*<0.05, \*\**p*<0.01, \*\*\**p*<0.001, \*\*\*\**p*<0.0001.

**Figure S4 (Related Figure 3).**
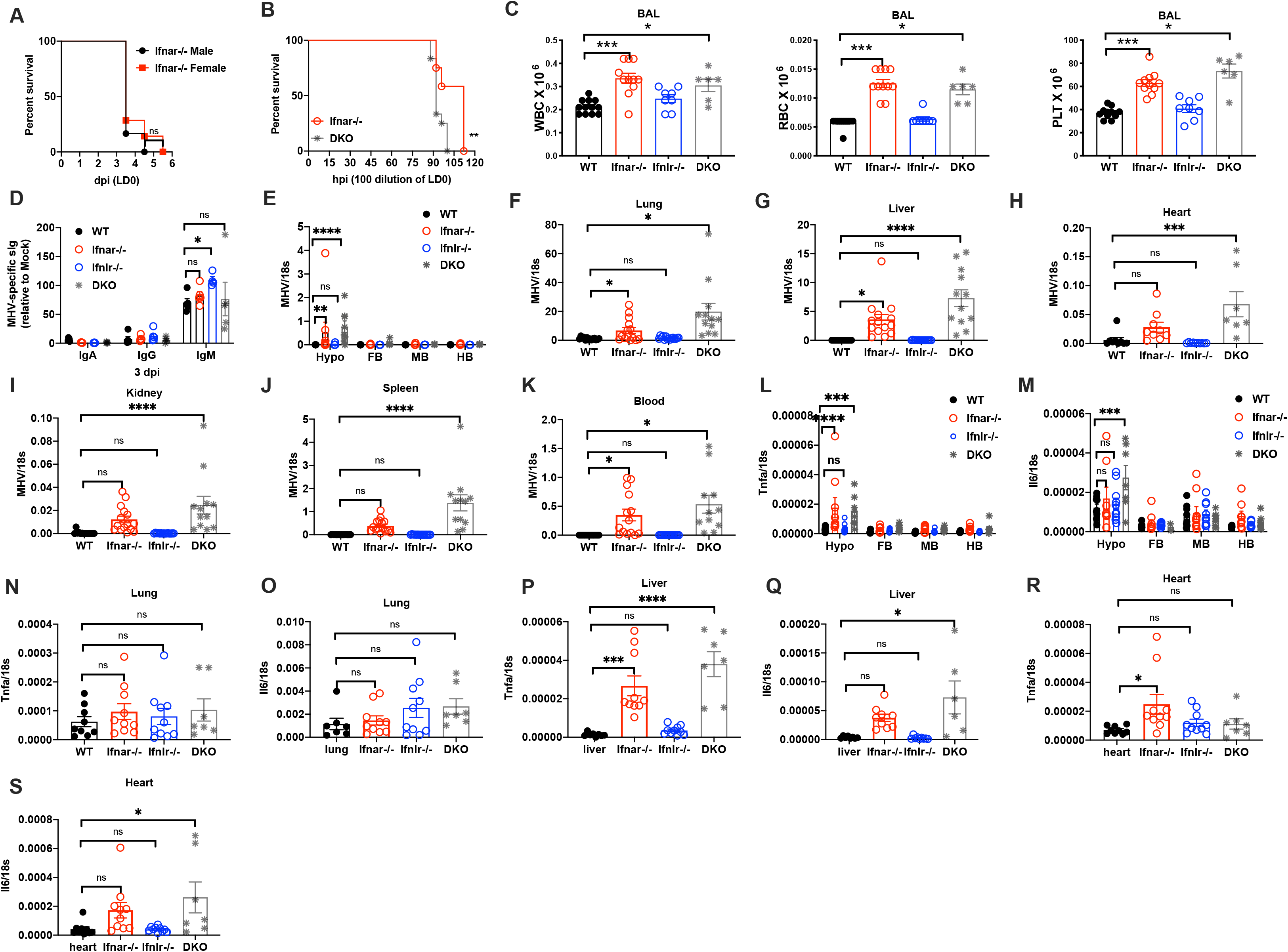
Absence of type I interferon response during respiratory MHV-A59 infection results in viremia and mortality, Related to Figure 3. **(A)** Survival rate of males comparing with females of IFNAR knockout (Ifnar1 −/−) mice. **(B)** Survival rate of Ifnar1 −/− comparing with double knockout of IFNAR and IFNLR (DKO) mice inoculated with MHV-A59 at 100 PFU (100 dilution of LD_0_). **(C)** white blood cells (WBC), red blood cells (RBC), and platelet (PLT) in bronchoalveolar lavage (BAL) from IFNAR (Ifnar−1/−), IFNLR (Ifnlr−/−), or double knockout (DKO), and wildtype (WT) mice infected with LD_0_ MHV-A59 on 3 dpi. **(D)** Serum levels of MHV-A59 specific IgA, IgG, and IgM from Ifnar1−/−, Ifnlr−/−, or DKO mice, and WT controls 3 days post respiratory MHV-A59 infection. **(E-K)** Transcriptional expression of MHV-A59, **(L-S)** Tnfa and Il6 in the indicated tissues from Ifnar1−/−, Ifnlr−/−, DKO, and WT mice infected with LD_0_ MHV-A59 on 3 dpi. Quantification of Data represented two independent experiment, 5-6 mice per group for every experiment. One-way analysis of variance (ANOVA) followed by Dunnett test for multiple groups comparation. Goups in (D,E, L, and M) without labeling indicates no significant difference. Results were presented as mean ± SEM. ns, not significant, \**p*<0.05, \*\**p*<0.01, \*\*\**p*<0.001, \*\*\*\**p*<0.0001.

**Figure S5 (Related to Figure 4).**
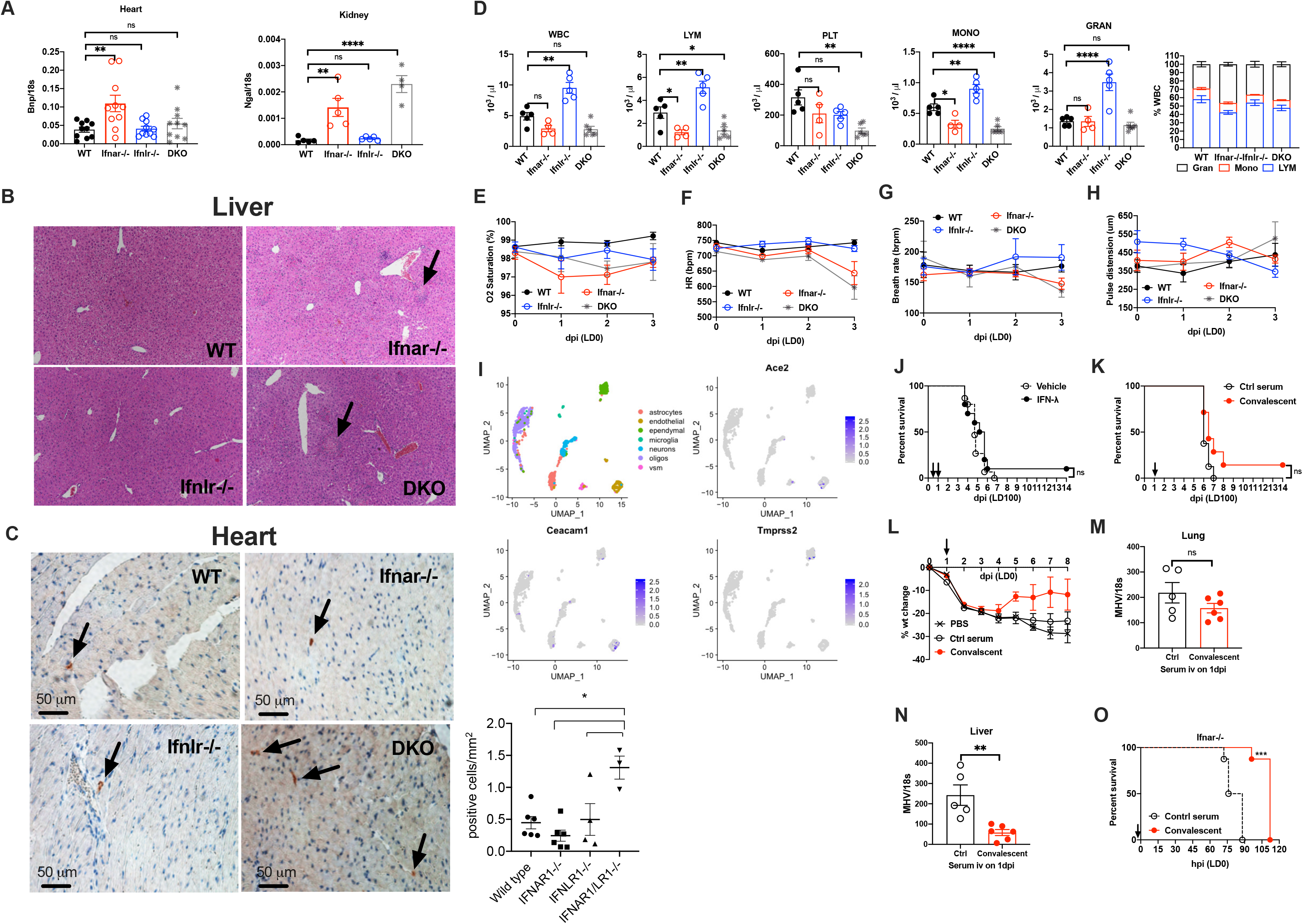
Deficiency of type I interferon response promotes brain invasion following MHV-A59 respiratory infection, Related to Figure 4. **(A)** Gene expression of brain natriuretic peptide (BNP), a marker of heart failure, and neutrophil gelatinase-associated lipocalin (NGAL), a marker for post-ischemic injury of kidney, in the heart and kidney respectively harvested on 3 days post MHV-A59 infection at LD_0_ of indicated knockout mice. **(B)** Representative images of H&E staining of the liver from indicated mice on 3dpi with LD_0_ MHV-A59 infection. **(C)** Representative images of F4/80 immunostaining of heart from same animals of (B). Positive staining is red, indicated by the arrows. Quantification of cells with positive staining was shown on the right. **(D)** Quantification of blood cells, **(E)** oxygen saturation, **(F)** heart rate, **(G)** breath rate, and **(H)** pulse distension from Ifnar1−/−, Ifnlr−/−, DKO, and WT mice infected with LD_0_ MHV-A59 on 3 dpi. **(I)** Cell clusters of mouse Ace2, Ceacam1, and Tmprss2 in hypothalamus from normal C57BL6/J mice using single-cell RNA sequencing analysis. **(J)** Survival rate of C57BL6/J mice intranasally inoculated with lethal MHV-A59 at LD_100_ followed by early interferon-λ (IFN-λ) treatment. Arrows indicated the timepoints (12 and 24 hours post infection) for IFN-λ intratracheal application. **(K)** Survival rate of C57BL6/J mice intranasally inoculated with lethal MHV-A59 at LD_100_ followed by convalescent serum treatment, normal serum as the control. Arrows indicated the timepoints (1 dpi) for intravenous serum injection. **(L)** Percentage of body weight change of C57BL6/J mice intranasally inoculated with unlethal MHV-A59 followed by convalescent serum treatment, normal serum and PBS as controls. Arrows indicated the timepoints (1 dpi) for intravenous injection of serums or PBS. Transcriptional expression of MHV-A59 in **(M)** the lung and **(N)** the liver on 3 dpi from C57BL6/J mice received convalescent or normal serum treatment 1 day after MHV-A59 respiratory infection. **(O)** Survival rate of Ifnar1−/− mice received convalescent serum intravenous injection right before intranasal MHV-A59 inoculation at LD_0_. Normal serum as the control. Arrows indicated the timepoint for intravenous serum injection. Data represented two independent experiment, 5-6 mice per group for every experiment. One-way analysis of variance (ANOVA) followed by Dunnett test for multiple groups comparation. Results were presented as mean ± SEM. ns, not significant, \**p*<0.05, \*\**p*<0.01, \*\*\**p*<0.001, \*\*\*\**p*<0.0001.

## EXPERIMENTAL MODEL AND SUBJECT DETAILS

### Mice

Young adult mice with age between 7 to 12 weeks were used. C57BL/6J, IFNAR1^−/−^, LysMCre, IFNAR^f/f^ animals were purchased from Jackson Laboratories. INFLR−/− were obtained from Sergei Kotenko. All animals were bred and maintained at Yale University. All animal experiments were performed according to institutional regulations upon review and approval of Yale University’s Institutional Animal Care and Use Committee.

Viral infections were performed in a biosafety level 2 (BSL2) facility and all experiments were done during the light period. Mice were intranasally inoculated with MHV-A59 following anesthesia with ketamine/xylazine cocktail. LD_50_ (Lethal Dose 50%) for every new stock of virus was tested in male young adult C57BL/6J mice, then PFU assay was applied using L2 cells infected with the same virus at ten-fold serial dilutions. In the current study, LD_50_ of MHV-A59 for young male C57BL/6J mice were between 10,000 and 160,000 PFU. All doses (LD_100_, LD_50_, and LD_0_) indicated here were referred to those tested from young male C57BL/6J mice. LD_50_ dose of MHV-A59 was uniformly applied to C57BL/6J mice except for the survival and rescue experiments; LD_0_ dose of MHV-A59 was applied to interferon deficiency mice as well as the wildtype controls because of their susceptibility to viral infection.

For rescue experiments, 1 μg interferon β or interferon λ (R & D Systems) in PBS with a total volume of 50 μl was intratracheally administrated to C57BL/6J mice challenged with lethal dose of MHV-A59 intranasally. Interferon treatments were applied at 12 and 24 hours or 36 and 48 hours post viral infection, respectively referring to the early or late course of disease. Convalescent serums were collected from survived mice at three weeks post MHV-A59 infection. Control serums were collected from normal healthy C57BL/6J mice. Serum treatments were applied through intravenous injection at 100ul per mouse 24 hours post viral infection for C57BL/6J mice or 10 minutes before viral infection for INFAR−/− mice respectively.

Vital signs, including blood oxygen saturation, breath rate, and heart rate post intranasal MHV-A59 inoculation were monitored via pulse oximetry using the MouseOx Plus (Starr Life Sciences Corp.). Core body temperature was measured by rectal probe thermometry (Physitemp TH-5 Thermalert).

### Virus

Recombinant murine coronavirus MHV-A59 were purchased from BEI resources (NR-43000), and were grown in BV2 cells. Viral titrations were tested using PFU assay in L2 cells infected with MHV-A59 at 10-fold serial dilutions.

## METHOD DETAILS

### Quantification of Plasma Cytokines

Plasma concentrations of IL-6, TNF-α and IL-1β were assayed by sandwich ELISA method. Recombinant, IL-6 (406-ML, R&D), TNF-α (410-MY, R&D) and IL-1β (401-ML-010, R&D) were utilized as standards with the highest concentration at 10ng/ml. Capture antibodies were anti-IL-6 (14-7061-85, eBioscience), anti-TNF-α (14-7423-85, eBioscience) and anti-IL-1β (14-7012-81, Invitrogen). Detection antibodies conjugated with biotin were anti-mouse-IL-6 (554402, BD Pharmingen), anti-mouse TNF-α (13-7349-81, Invitrogen) and anti-mouse IL-1β (13-7112-85, eBioscience). Following the incubation with HRP-conjugated streptavidin (554066, BD Biosciences) and TMB substrate reagent (555214, BD Biosciences), color of the plates were closely monitored every minute and plates were read at 450nm instantly after the addition of stop solution (1N HCL). Interferon-a and Interferon-β measurements were performed per protocols from manufacturers.

### Quantification of Serum MHV-A59 specific immunoglobulins

Heat inactivated sera were assayed for immunoglobulins by ELISA in plates coated with 4^5 PFU/well of MHV-A59 viruses that were diluted in 100ul sodium carbonate buffer (PH 8.0). 1ul serum for IgGs and IgM measurements, or 5ul serum for IgA measurement diluted in PBS with 1%BSA to a total volume of 100ul per well were incubate at 4°C overnight. Plates were afterwards incubated with HRP-conjugated secondary goat anti-mouse IgGs, IgM, or IgA antibodies (abcam) with 10,000 dilution in PBS with 1% BSA. TMB substrate reagent and stop solution were the same as those for cytokine detection. Plates were read at 450nm and 570nm.

### Quantification of Organ Injury Markers and Hematology Analysis

Cardiac Troponin-I (CTNI) concentration and Alanine Aminotransferase (ALT) activity in the blood were measured by kits per manufacturers’ instructions (Life Diagnostics and Cayman Chemical, respectively). 50ul of whole blood per mouse were collected via retro-orbital bleeding in heparin coated tubes then counted immediately using HemaTrue (HESK).

### Virus Titration

Virus titration was analyzed as previous described (Leibowitz et al., 2011). Briefly, 90% confluent L2 cells were infected with 10-fold serial dilutions of MHV-A59 virus. After one-hour incubation, each well was overlaid with 1:1 mixture of growth median and 1.2% Avicel for four days. Cells were then fixed and stained with crystal violet, and plaques were enumerated in triplicated.

### Quantification of Viral Load

Viral load in the tissue were quantified using q-PCR, PFU or TCID50 in the current study. Primers to amplify nucleocapsid protein of MHV-A59 were used to quantify viral RNA. Viral load were presented as the relative expression comparing with housekeeping gene 18s. Upper left pulmonary lobe was uniformly applied for PFU assay. 10-fold serial dilutions of homogenized lung tissue were incubated with L2 cells, then plaque stain was performed after 48 hours incubation. TCID50 were performed by infecting serial dilutions of BAL samples onto L2 monolayers and incubated for 48 hours. Cells were then stained with crystal violet to determine the dilution that is sufficient to kill the cells.

### qRT-PCR

Tissues were homogenized in 1ml RNA-Bee (Tel-Test, Inc) using a FastPrep-24 5G homogenizer (MP Biomedicals). RNA was purified using QIAGEN RNeasy columns according to the manufacturer’s instructions. cDNA was generated with reverse transcriptase (Clontech) using oligo-dT6 primers (Sigma-Aldrich). qRT-PCR was performed on a CFX96 Real-Time System (Bio-Rad) using PerfeCTa SYBR Green SuperMix (Quanta Biosciences). Relative expression units were calculated as transcript levels of target genes relative to *18s*. Primers used for qRT-PCR are listed in Table S1.

### Immunohistochemistry

Mice were euthanized and perfused with PBS or fixative. All tissues were afterwards immersion-fixed in 10% neutral buffered formalin. Hematoxylin and Eosin (H&E), or Martius, Scarlet and Blue (MSB) staining was applied, then slides were assessed by a blinded pathologist.

### Single-cell RNA sequencing

Sequence data was obtained from Gene Expression Omnibus, accession code GSE74672 (Romanov et al., 2017). Expression matrix combined with metadata were loaded into Seurat v3.0 (Butler et al., 2018). Feature distribution and filtering (nFeature_RNA > 200 & nFeature_RNA < 2500) were performed with FeatureScatter and subset functions. Then, data normalization, assignment of variable genes and scaling were performed with Seurat’s NormalizeData, FindVariableFeatures and ScaleData functions, respectively. Dimensionality reduction was performed with RunPCA function and significant dimensions were evaluated with ElbowPlot and JackStrawPlot. Clustering of cells was performed with Seurat’s FindNeighbors (dims=1:30) and FindClusters (resolution=0.8) functions, while optimal cluster resolution was determined using clustree (Zappia and Oshlack, 2018). Finally, UMAP was performed with Seurat’s RunUMAP function (dims =1:30) and gene expression of features of interest were visualized with FeaturePlot function.

## QUANTIFICATION AND STATISTICAL ANALYSIS

Statistical analyses were performed using Prism 8.0 (GraphPad Software, Inc.). Student’s *t* test was used for two groups comparison. More than two groups were compared using one-way analysis of variance (ANOVA) followed by Dunnett test. Samples at different time points from multiple groups were analyzed using two-way ANOVA followed by Tukey test. A p value less than 0.05 was considered statistically significant. Data are presented as the mean ± SEM. * p<0.05, ** p<0.01, *** p<0.001, **** p<0.0001

**Table S1.**
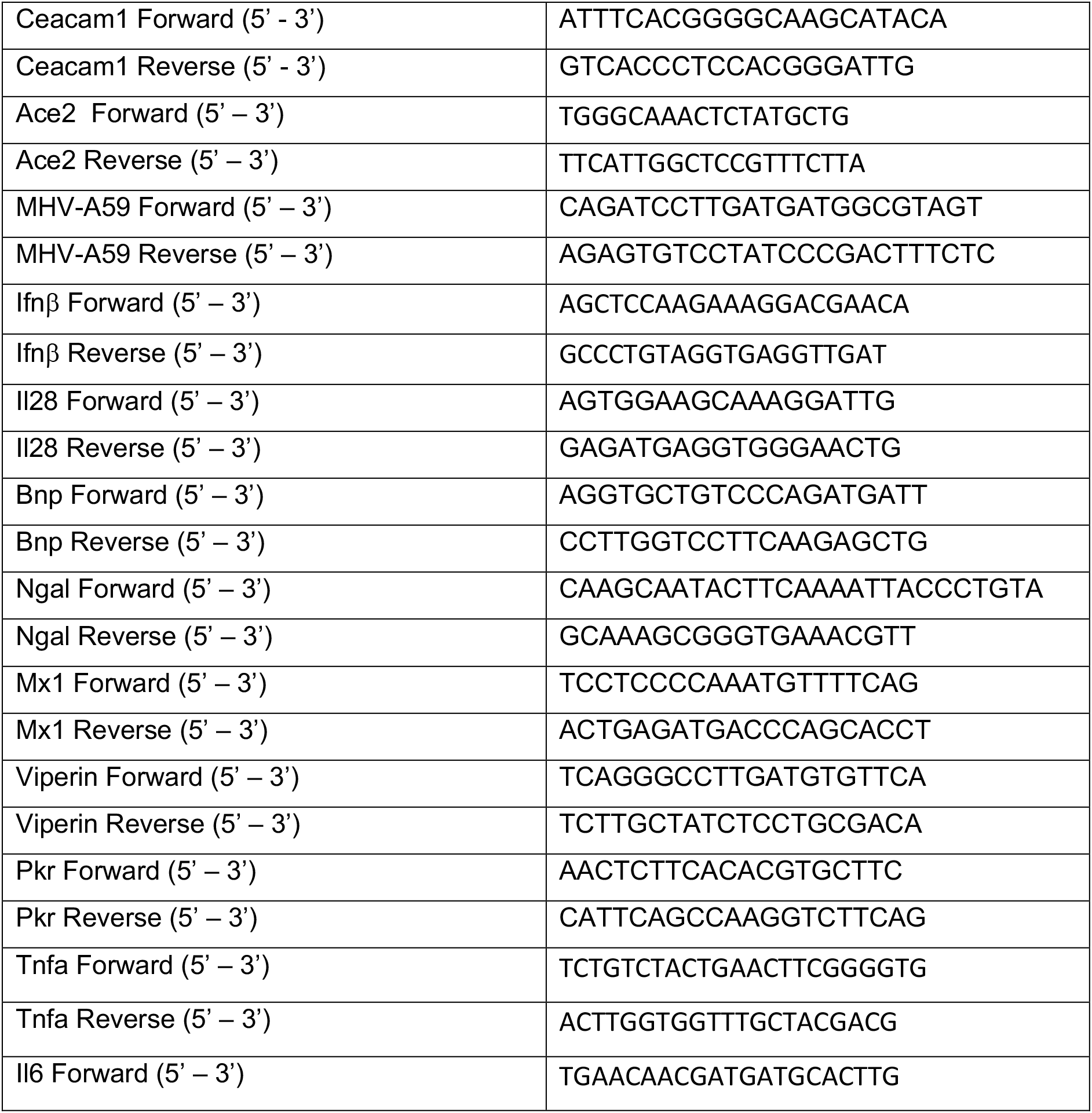
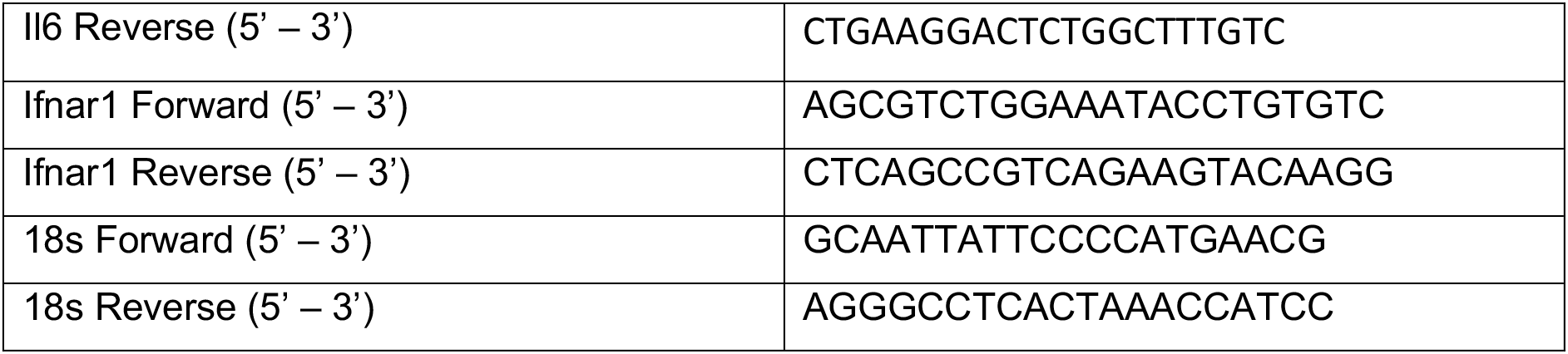
Primers for qRT-PCR, Related to Key Resources Table

## References

Bao, L., Deng, W., Huang, B., Gao, H., Liu, J., Ren, L., Wei, Q., Yu, P., Xu, Y., Qi, F., et al. (2020). The pathogenicity of SARS-CoV-2 in hACE2 transgenic mice. Nature 583, 830–833.

Butler, A., Hoffman, P., Smibert, P., Papalexi, E., and Satija, R. (2018). Integrating single-cell transcriptomic data across different conditions, technologies, and species. Nat Biotechnol 36, 411–420.

Cervantes-Barragan, L., Kalinke, U., Zust, R., Konig, M., Reizis, B., Lopez-Macias, C., Thiel, V., and Ludewig, B. (2009). Type I IFN-mediated protection of macrophages and dendritic cells secures control of murine coronavirus infection. J Immunol 182, 1099–1106.

Chan, J.F., Zhang, A.J., Yuan, S., Poon, V.K., Chan, C.C., Lee, A.C., Chan, W.M., Fan, Z., Tsoi, H.W., Wen, L., et al. (2020). Simulation of the clinical and pathological manifestations of Coronavirus Disease 2019 (COVID-19) in golden Syrian hamster model: implications for disease pathogenesis and transmissibility. Clin Infect Dis.

Compton, S.R., Ball-Goodrich, L.J., Johnson, L.K., Johnson, E.A., Paturzo, F.X., and Macy, J.D. (2004). Pathogenesis of enterotropic mouse hepatitis virus in immunocompetent and immunodeficient mice. Comp Med 54, 681–689.

Compton, S.R., Stephensen, C.B., Snyder, S.W., Weismiller, D.G., and Holmes, K.V. (1992). Coronavirus species specificity: murine coronavirus binds to a mouse-specific epitope on its carcinoembryonic antigen-related receptor glycoprotein. J Virol 66, 7420–7428.

Coronaviridae Study Group of the International Committee on Taxonomy of, V. (2020). The species Severe acute respiratory syndrome-related coronavirus: classifying 2019-nCoV and naming it SARS-CoV-2. Nat Microbiol 5, 536–544.

DeAlbuquerque, N., Baig, E., Xuezhong, M., Shalev, I., Phillips, M.J., Habal, M., Leibowitz, J., McGilvray, I., Butany, J., Fish, E., et al. (2006). Murine hepatitis virus strain 1 as a model for severe acute respiratory distress syndrome (SARS). Adv Exp Med Biol 581, 373–378.

DeBiasi, R.L., Song, X., Delaney, M., Bell, M., Smith, K., Pershad, J., Ansusinha, E., Hahn, A., Hamdy, R., Harik, N., et al. (2020). Severe Coronavirus Disease-2019 in Children and Young Adults in the Washington, DC, Metropolitan Region. J Pediatr 223, 199–203 e191.

Dery, K.J., Kujawski, M., Grunert, D., Wu, X., Ngyuen, T., Cheung, C., Yim, J.H., and Shively, J.E. (2014). IRF-1 regulates alternative mRNA splicing of carcinoembryonic antigen-related cell adhesion molecule 1 (CEACAM1) in breast epithelial cells generating an immunoreceptor tyrosine-based inhibition motif (ITIM) containing isoform. Mol Cancer 13, 64.

Ergun, S., Kilik, N., Ziegeler, G., Hansen, A., Nollau, P., Gotze, J., Wurmbach, J.H., Horst, A., Weil, J., Fernando, M., et al. (2000). CEA-related cell adhesion molecule 1: a potent angiogenic factor and a major effector of vascular endothelial growth factor. Mol Cell 5, 311–320.

Farmer, D.G., Dutschmann, M., Paton, J.F., Pickering, A.E., and McAllen, R.M. (2016). Brainstem sources of cardiac vagal tone and respiratory sinus arrhythmia. J Physiol 594, 7249–7265.

Frieman, M., and Baric, R. (2008). Mechanisms of severe acute respiratory syndrome pathogenesis and innate immunomodulation. Microbiol Mol Biol Rev 72, 672–685, Table of Contents.

Godfraind, C., Langreth, S.G., Cardellichio, C.B., Knobler, R., Coutelier, J.P., Dubois-Dalcq, M., and Holmes, K.V. (1995). Tissue and cellular distribution of an adhesion molecule in the carcinoembryonic antigen family that serves as a receptor for mouse hepatitis virus. Lab Invest 73, 615–627.

Gorbalenya, A.E., Snijder, E.J., and Spaan, W.J. (2004). Severe acute respiratory syndrome coronavirus phylogeny: toward consensus. J Virol 78, 7863–7866.

Grajales-Reyes, G.E.C., Marco (2020). Interferon responses in viral pneumonias. Science 369, 626–627.

Grasselli, G., Greco, M., Zanella, A., Albano, G., Antonelli, M., Bellani, G., Bonanomi, E., Cabrini, L., Carlesso, E., Castelli, G., et al. (2020). Risk Factors Associated With Mortality Among Patients With COVID-19 in Intensive Care Units in Lombardy, Italy. JAMA Intern Med.

Gu, H., Chen, Q., Yang, G., He, L., Fan, H., Deng, Y.Q., Wang, Y., Teng, Y., Zhao, Z., Cui, Y., et al. (2020). Adaptation of SARS-CoV-2 in BALB/c mice for testing vaccine efficacy. Science.

Gupta, A., Madhavan, M.V., Sehgal, K., Nair, N., Mahajan, S., Sehrawat, T.S., Bikdeli, B., Ahluwalia, N., Ausiello, J.C., Wan, E.Y., et al. (2020). Extrapulmonary manifestations of COVID-19. Nat Med 26, 1017–1032.

Hadjadj, J., Yatim, N., Barnabei, L., Corneau, A., Boussier, J., Smith, N., Pere, H., Charbit, B., Bondet, V., Chenevier-Gobeaux, C., et al. (2020). Impaired type I interferon activity and inflammatory responses in severe COVID-19 patients. Science 369, 718–724.

Hamming, I., Timens, W., Bulthuis, M.L., Lely, A.T., Navis, G., and van Goor, H. (2004). Tissue distribution of ACE2 protein, the functional receptor for SARS coronavirus. A first step in understanding SARS pathogenesis. J Pathol 203, 631–637.

Hassan, A.O., Case, J.B., Winkler, E.S., Thackray, L.B., Kafai, N.M., Bailey, A.L., McCune, B.T., Fox, J.M., Chen, R.E., Alsoussi, W.B., et al. (2020). A SARS-CoV-2 Infection Model in Mice Demonstrates Protection by Neutralizing Antibodies. Cell 182, 744–753 e744.

Hundt, M.A., Deng, Y., Ciarleglio, M.M., Nathanson, M.H., and Lim, J.K. (2020). Abnormal Liver Tests in COVID-19: A Retrospective Observational Cohort Study of 1827 Patients in a Major U.S. Hospital Network. Hepatology.

Ing, E.B., Xu, Q.A., Salimi, A., and Torun, N. (2020). Physician deaths from corona virus (COVID-19) disease. Occup Med (Lond) 70, 370–374.

Israelow, B., Song, E., Mao, T., Lu, P., Meir, A., Liu, F., Alfajaro, M.M., Wei, J., Dong, H., Homer, R.J., et al. (2020). Mouse model of SARS-CoV-2 reveals inflammatory role of type I interferon signaling. J Exp Med 217.

Jiang, R.D., Liu, M.Q., Chen, Y., Shan, C., Zhou, Y.W., Shen, X.R., Li, Q., Zhang, L., Zhu, Y., Si, H.R., et al. (2020). Pathogenesis of SARS-CoV-2 in Transgenic Mice Expressing Human Angiotensin-Converting Enzyme 2. Cell 182, 50–58 e58.

Kim, Y.I., Kim, S.G., Kim, S.M., Kim, E.H., Park, S.J., Yu, K.M., Chang, J.H., Kim, E.J., Lee, S., Casel, M.A.B., et al. (2020). Infection and Rapid Transmission of SARS-CoV-2 in Ferrets. Cell Host Microbe 27, 704–709 e702.

Klein, S.L., Hodgson, A., and Robinson, D.P. (2012). Mechanisms of sex disparities in influenza pathogenesis. J Leukoc Biol 92, 67–73.

Leibowitz, J., Kaufman, G., and Liu, P. (2011). Coronaviruses: propagation, quantification, storage, and construction of recombinant mouse hepatitis virus. Curr Protoc Microbiol Chapter 15, Unit 15E 11.

Leibowitz, J.L., Srinivasa, R., Williamson, S.T., Chua, M.M., Liu, M., Wu, S., Kang, H., Ma, X.Z., Zhang, J., Shalev, I., et al. (2010). Genetic determinants of mouse hepatitis virus strain 1 pneumovirulence. J Virol 84, 9278–9291.

Li, K., Wohlford-Lenane, C.L., Channappanavar, R., Park, J.E., Earnest, J.T., Bair, T.B., Bates, A.M., Brogden, K.A., Flaherty, H.A., Gallagher, T., et al. (2017). Mouse-adapted MERS coronavirus causes lethal lung disease in human DPP4 knockin mice. Proc Natl Acad Sci U S A 114, E3119–E3128.

Lucas, C., Wong, P., Klein, J., Castro, T.B.R., Silva, J., Sundaram, M., Ellingson, M.K., Mao, T., Oh, J.E., Israelow, B., et al. (2020). Longitudinal analyses reveal immunological misfiring in severe COVID-19. Nature 584, 463–469.

Mallapaty, S. (2020). The coronavirus is most deadly if you are older and male - new data reveal the risks. Nature.

McCray, P.B., Jr., Pewe, L., Wohlford-Lenane, C., Hickey, M., Manzel, L., Shi, L., Netland, J., Jia, H.P., Halabi, C., Sigmund, C.D., et al. (2007). Lethal infection of K18-hACE2 mice infected with severe acute respiratory syndrome coronavirus. J Virol 81, 813–821.

Miura, T.A., Travanty, E.A., Oko, L., Bielefeldt-Ohmann, H., Weiss, S.R., Beauchemin, N., and Holmes, K.V. (2008). The spike glycoprotein of murine coronavirus MHV-JHM mediates receptor-independent infection and spread in the central nervous systems of Ceacam1a−/− Mice. J Virol 82, 755–763.

Momtazmanesh, S., Shobeiri, P., Hanaei, S., Mahmoud-Elsayed, H., Dalvi, B., and Malakan Rad, E. (2020). Cardiovascular disease in COVID-19: a systematic review and meta-analysis of 10,898 patients and proposal of a triage risk stratification tool. Egypt Heart J 72, 41.

Montalvan, V., Lee, J., Bueso, T., De Toledo, J., and Rivas, K. (2020). Neurological manifestations of COVID-19 and other coronavirus infections: A systematic review. Clin Neurol Neurosurg 194, 105921.

Munster, V.J., Feldmann, F., Williamson, B.N., van Doremalen, N., Perez-Perez, L., Schulz, J., Meade-White, K., Okumura, A., Callison, J., Brumbaugh, B., et al. (2020). Respiratory disease in rhesus macaques inoculated with SARS-CoV-2. Nature.

Naqvi, A.A.T., Fatima, K., Mohammad, T., Fatima, U., Singh, I.K., Singh, A., Atif, S.M., Hariprasad, G., Hasan, G.M., and Hassan, M.I. (2020). Insights into SARS-CoV-2 genome, structure, evolution, pathogenesis and therapies: Structural genomics approach. Biochim Biophys Acta Mol Basis Dis 1866, 165878.

Nguyen, L.H., Drew, D.A., Graham, M.S., Joshi, A.D., Guo, C.G., Ma, W., Mehta, R.S., Warner, E.T., Sikavi, D.R., Lo, C.H., et al. (2020). Risk of COVID-19 among front-line health-care workers and the general community: a prospective cohort study. Lancet Public Health.

Park, A., and Iwasaki, A. (2020). Type I and Type III Interferons - Induction, Signaling, Evasion, and Application to Combat COVID-19. Cell Host Microbe 27, 870–878.

Perlman, S., Jacobsen, G., and Afifi, A. (1989). Spread of a neurotropic murine coronavirus into the CNS via the trigeminal and olfactory nerves. Virology 170, 556–560.

Phillips, J.M., and Weiss, S.R. (2011). Pathogenesis of neurotropic murine coronavirus is multifactorial. Trends Pharmacol Sci 32, 2–7.

Pinol, R.A., Zahler, S.H., Li, C., Saha, A., Tan, B.K., Skop, V., Gavrilova, O., Xiao, C., Krashes, M.J., and Reitman, M.L. (2018). Brs3 neurons in the mouse dorsomedial hypothalamus regulate body temperature, energy expenditure, and heart rate, but not food intake. Nat Neurosci 21, 1530–1540.

Puelles, V.G., Lutgehetmann, M., Lindenmeyer, M.T., Sperhake, J.P., Wong, M.N., Allweiss, L., Chilla, S., Heinemann, A., Wanner, N., Liu, S., et al. (2020). Multiorgan and Renal Tropism of SARS-CoV-2. N Engl J Med 383, 590–592.

Ragab, D., Salah Eldin, H., Taeimah, M., Khattab, R., and Salem, R. (2020). The COVID-19 Cytokine Storm; What We Know So Far. Front Immunol 11, 1446.

Romanov, R.A., Zeisel, A., Bakker, J., Girach, F., Hellysaz, A., Tomer, R., Alpar, A., Mulder, J., Clotman, F., Keimpema, E., et al. (2017). Molecular interrogation of hypothalamic organization reveals distinct dopamine neuronal subtypes. Nat Neurosci 20, 176–188.

Romero-Sanchez, C.M., Diaz-Maroto, I., Fernandez-Diaz, E., Sanchez-Larsen, A., Layos-Romero, A., Garcia-Garcia, J., Gonzalez, E., Redondo-Penas, I., Perona-Moratalla, A.B., Del Valle-Perez, J.A., et al. (2020). Neurologic manifestations in hospitalized patients with COVID-19: The ALBACOVID registry. Neurology 95, e1060–e1070.

Sun, J., Zhuang, Z., Zheng, J., Li, K., Wong, R.L., Liu, D., Huang, J., He, J., Zhu, A., Zhao, J., et al. (2020a). Generation of a Broadly Useful Model for COVID-19 Pathogenesis, Vaccination, and Treatment. Cell 182, 734–743 e735.

Sun, S.H., Chen, Q., Gu, H.J., Yang, G., Wang, Y.X., Huang, X.Y., Liu, S.S., Zhang, N.N., Li, X.F., Xiong, R., et al. (2020b). A Mouse Model of SARS-CoV-2 Infection and Pathogenesis. Cell Host Microbe 28, 124–133 e124.

Takahashi, T., Ellingson, M.K., Wong, P., Israelow, B., Lucas, C., Klein, J., Silva, J., Mao, T., Oh, J.E., Tokuyama, M., et al. (2020). Sex differences in immune responses that underlie COVID-19 disease outcomes. Nature.

Walz, L., Cohen, A.J., Rebaza, A.P., Vanchieri, J., Slade, M.D., Dela Cruz, C.S., and Sharma, L. (2020). Janus Kinase-Inhibitor and Type I Interferon Ability to Produce Favorable Clinical Outcomes in COVID-19 Patients: A Systematic Review and Meta-Analysis. medRxiv.

Williamson, E.J., Walker, A.J., Bhaskaran, K., Bacon, S., Bates, C., Morton, C.E., Curtis, H.J., Mehrkar, A., Evans, D., Inglesby, P., et al. (2020). Factors associated with COVID-19-related death using OpenSAFELY. Nature 584, 430–436.

Yang, Z., Du, J., Chen, G., Zhao, J., Yang, X., Su, L., Cheng, G., and Tang, H. (2014). Coronavirus MHV-A59 infects the lung and causes severe pneumonia in C57BL/6 mice. Virol Sin 29, 393–402.

Zappia, L., and Oshlack, A. (2018). Clustering trees: a visualization for evaluating clusterings at multiple resolutions. Gigascience 7.

Zhao, Q., Meng, M., Kumar, R., Wu, Y., Huang, J., Deng, Y., Weng, Z., and Yang, L. (2020). Lymphopenia is associated with severe coronavirus disease 2019 (COVID-19) infections: A systemic review and meta-analysis. Int J Infect Dis 96, 131–135.

Zheng, K.I., Feng, G., Liu, W.Y., Targher, G., Byrne, C.D., and Zheng, M.H. (2020). Extrapulmonary complications of COVID-19: A multisystem disease? J Med Virol.

Ziegler, C.G.K., Allon, S.J., Nyquist, S.K., Mbano, I.M., Miao, V.N., Tzouanas, C.N., Cao, Y., Yousif, A.S., Bals, J., Hauser, B.M., et al. (2020). SARS-CoV-2 Receptor ACE2 Is an Interferon-Stimulated Gene in Human Airway Epithelial Cells and Is Detected in Specific Cell Subsets across Tissues. Cell 181, 1016–1035 e1019.

